# CTCF-mediated insulation and chromatin environment modulate *Car5b* escape from X inactivation

**DOI:** 10.1101/2023.05.04.539469

**Authors:** He Fang, Ana R. Tronco, Giancarlo Bonora, Truong Nguyen, Jitendra Thakur, Joel B. Berletch, Galina N. Filippova, Steven Henikoff, Jay Shendure, William S. Noble, Christine M. Disteche, Xinxian Deng

## Abstract

**Background:** The number and escape levels of genes that escape X chromosome inactivation (XCI) in female somatic cells vary among tissues and cell types, potentially contributing to specific sex differences. Here we investigate the role of CTCF, a master chromatin conformation regulator, in regulating escape from XCI. CTCF binding profiles and epigenetic features were systematically examined at constitutive and facultative escape genes using mouse allelic systems to distinguish the inactive X (Xi) and active X (Xa) chromosomes.

**Results:** We found that escape genes are located inside domains flanked by convergent arrays of CTCF binding sites, consistent with the formation of loops. In addition, strong and divergent CTCF binding sites often located at the boundaries between escape genes and adjacent neighbors subject to XCI would help insulate domains. Facultative escapees show clear differences in CTCF binding dependent on their XCI status in specific cell types/tissues. Concordantly, deletion but not inversion of a CTCF binding site at the boundary between the facultative escape gene *Car5b* and its silent neighbor *Siah1b* resulted in loss of *Car5b* escape. Reduced CTCF binding and enrichment of a repressive mark over *Car5b* in cells with a boundary deletion indicated loss of looping and insulation. In mutant lines in which either the Xi-specific compact structure or its H3K27me3 enrichment was disrupted, escape genes showed an increase in gene expression and associated active marks, supporting the roles of the 3D Xi structure and heterochromatic marks in constraining levels of escape.

**Conclusion:** Our findings indicate that escape from XCI is modulated both by looping and insulation of chromatin via convergent arrays of CTCF binding sites and by compaction and epigenetic features of the surrounding heterochromatin.

## Background

X chromosome inactivation (XCI) silences one of the two X chromosomes in mammalian females to equalize X-linked gene expression between males (XY) and females (XX) [1]. XCI is a chromosome-wide epigenetic process orchestrated by the long non-coding RNA (lncRNA) *Xist* (X-inactive specific transcript) during early female embryogenesis [2,3]. *Xist* RNA cis-coats the future inactive X chromosome (Xi), and recruits or removes specific proteins and chromatin modifications to silence genes. As a result, the Xi acquires a specific pattern of epigenetic hallmarks due to depletion of active histone modifications and enrichment in repressive modifications. Xi-specific features also include enrichment of the histone variant macroH2A, hyper-methylation of CpG islands, and late DNA replication. In addition, the Xi becomes highly condensed and is reshaped into a heterochromatic bipartite structure containing two compact superdomains of long-range chromatin interactions separated by a boundary at the conserved macrosatellite repeat region *Dxz4*. The multi-layer epigenetic regulation of the Xi ensures that its silent state is stably maintained in somatic cells. However, some genes escape XCI [4,5], including about 15-30% of human and 3-7% of mouse X-linked genes, resulting in higher expression levels in females compared to males, which potentially contributes to phenotypic sex differences. The number and levels of genes that escape XCI vary among species, tissues, and cell types [6–10]. Some genes (“constitutive escapees”) escape XCI in a ubiquitous manner, while others (“facultative escapees”) only escape in certain cell types and tissues, resulting in specific sex differences.

The mechanisms by which escapees evade the multiple layers of heterochromatin control inherent to XCI and the factors that regulate expression levels from the Xi in specific cell types, tissues, and species are not well understood. A better understanding of the mechanisms of escape and the factors that influence it would help define the roles of escapees in sex differences. Emerging evidence has shown that an important master regulator of chromatin structure, CTCF (CCCTC-binding factor), may play an important role in regulating escape from XCI. We previously reported that CTCF binds to the 5’ end of two mouse constitutive escapees, *Kdm5c* and *Eif2s3x,* where it might function as a boundary to insulate these genes from neighbors that are subject to XCI [11]. In support of this model, CTCF profiles we obtained by allele-specific ChIP-seq in two systems with skewed XCI, a fibroblast cell line (Patski) and adult tissues from F1 hybrid mice, have clearly shown that CTCF binding is reduced on the Xi versus the Xa (active X), except at escape regions [8]. Interestingly, *Car5b* that escapes XCI in Patski cells but not in mouse brain is flanked by CTCF binding sites in Patski cells, supporting the role of CTCF in regulation of escape. Furthermore, it has been reported that BAC transgenes containing *Kdm5c* and its distal (3’ end) boundary inserted at random on the Xi maintain their escape status at various insertion sites during mouse embryonic stem cell differentiation [12]. However, in the absence of the distal boundary there is inappropriate spreading of escape from the *Kdm5c* BAC to neighbor genes normally subject to XCI located at the 3’ end of the insertion site, supporting the idea that insulator elements separate domains of escape from silenced domains. An *in vivo* transgenic mouse study further showed maintenance of escape after insertion of BACs containing either a mouse (*Kdm5c*) or a human (*RPS4X,* ribosomal protein S4 X-linked) escape gene and their flanking neighbor genes into the *Hprt* (hypoxanthine phosphoribosyltransferase) gene, which is subject to XCI [13]. More intriguingly, such insulation can be achieved by artificial CTCF tethering via CRISPR/dCas, which has been recently reported to help reactivate *MECP2* on the Xi when combined with CpG demethylation [14]. While these findings strongly suggest that boundaries or other *cis*-elements regulate escape, the molecular mechanisms remain elusive.

In this study we systematically examined allele-specific CTCF binding patterns and epigenetic features at constitutive and variable escapees in Patski cells and in adult mouse F1 hybrid tissues, using ChIP-seq, CUT&RUN, and ATAC-seq analyses. We found that escapees on the Xi are often located inside domains of several hundred kilobases, flanked by convergent arrays of CTCF binding sites. In addition, strong and divergent CTCF binding sites are often located at the boundaries between escape genes and adjacent genes subject to XCI. To functionally test the role of CTCF binding in escape, we applied an allele-specific CRISPR/Cas9 approach to delete or invert a strong CTCF binding site at the boundary proximal to the promoter of the variable escape gene *Car5b* on the Xi or Xa in Patski cells. We also explored the effects of disrupting condensation and heterochromatic environment of the entire Xi on escape. These studies affirm the roles of boundary elements such as CTCF, and of histone modifications and chromatin condensation in modulation of escape.

## Results

### Distribution of CTCF binding around escape genes

We systematically examined epigenetic features around escape genes in interspecific mouse systems with skewed XCI in which Xi and Xa alleles can be identified based on species-specific SNPs. These systems include the embryonic fibroblast line Patski and brain from F1 hybrid female mice, in which we have previously reported the inactivation status of genes and the overall distribution of CTCF binding obtained by ChIP-seq [8,15]. Detailed examination of CTCF binding sites was done here to identify Xi-peaks with strong CTCF motif scores and convergent orientations at regions containing six constitutive (*Ddx3x*, *Kdm6a*, *Eif2s3x*, *Xist*, *Pbdc1*, and *Kdm5c*) and two tissue-specific facultative escapees (*Shroom4* and *Car5b*) (Fig. 1A, B, Fig. 2A-F, Additional file 2: Table S1).

**Figure 1.**
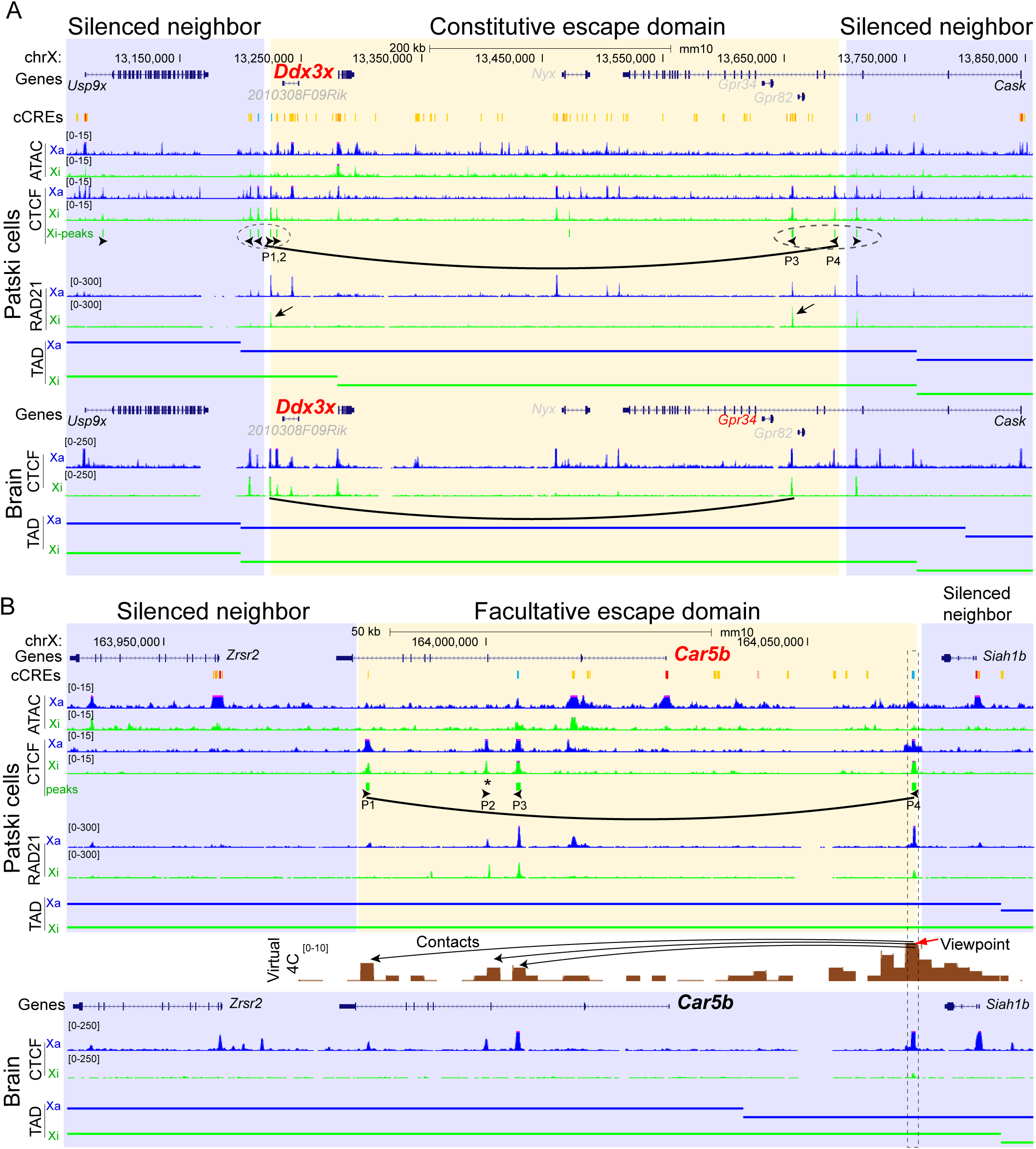
Escape domains at the constitutive escape gene *Ddx3x* and facultative escape gene *Car5b*. **A**. UCSC browser view of the constitutive *Ddx3x* domain in Patski cells and in mouse brain. Allelic profiles of ATAC-seq reads, CTCF and RAD21 ChIP-seq reads and TADs on the Xa (blue) and Xi (green are shown). The boundaries between the putative escape domain (yellow) and its flanking silent regions (blue) are marked by several CTCF binding sites with divergent orientation (circled black arrowheads). Inside of the escape domain convergent arrays of CTCF binding sites and RAD21 peaks suggest loop interactions (curve line) mediated by forward motifs P1, P2, and reverse motifs P3, P4 (black arrowheads). Only CTCF Xi-peaks with a conserved motif score ≥10 from CTCFBSDB are marked. The *Ddx3x* escape domain approximately overlaps with a Xi TAD. ENCODE candidate *cis*-regulatory elements (cCREs) combined for all mouse cell types are shown for promoters in red, enhancers in yellow, and conserved CTCF sites in cyan. Profiles of CTCF ChIP-seq reads and TADs in mouse brain are shown. Note that *Gpr34* which is not expressed in Patski cells, escapes XCI in brain (Additional file1: Fig. S1B). Genes known to escape XCI are marked in red, genes subject to XCI, in black, and genes that are not expressed or not assessable due to lack of SNPs, in grey. **B**. Same analysis for the facultative escape *Car5b* in Patski cells and mouse brain. CTCF binding site P4 is located between *Car5b* and its inactivated neighbor *Siah1b*. Three CTCF binding sites are located within the body of *Car5b* (P1, P2 and P3). While no CTCF peak was called at P2 on the Xi using our cutoff for peak calling (star), CTCF binding regions P1 and P2 include strong CTCF forward motifs, and P2 also binds to RAD21 on the Xi, suggesting important anchor sites. In support of this, the virtual 4C contact plot derived from Hi-C data at 1kb resolution using a 5kb window around CTCF peak P4 (marked by red arrow) as a viewpoint reveals contacts that overlap peaks P1, P2 and P3, suggesting looping events involving three regions within the body of *Car5b*. There is no correlation with the TAD structure. Note that the *Car5b* promoter and enhancer marked by ATAC-seq peaks and ENCODE cCREs (red and yellow bars) lack strong CTCF peaks on the Xi. Importantly, there is no or very little CTCF binding on the Xi in brain where *Car5b* does not escape XCI.

**Figure 2.**
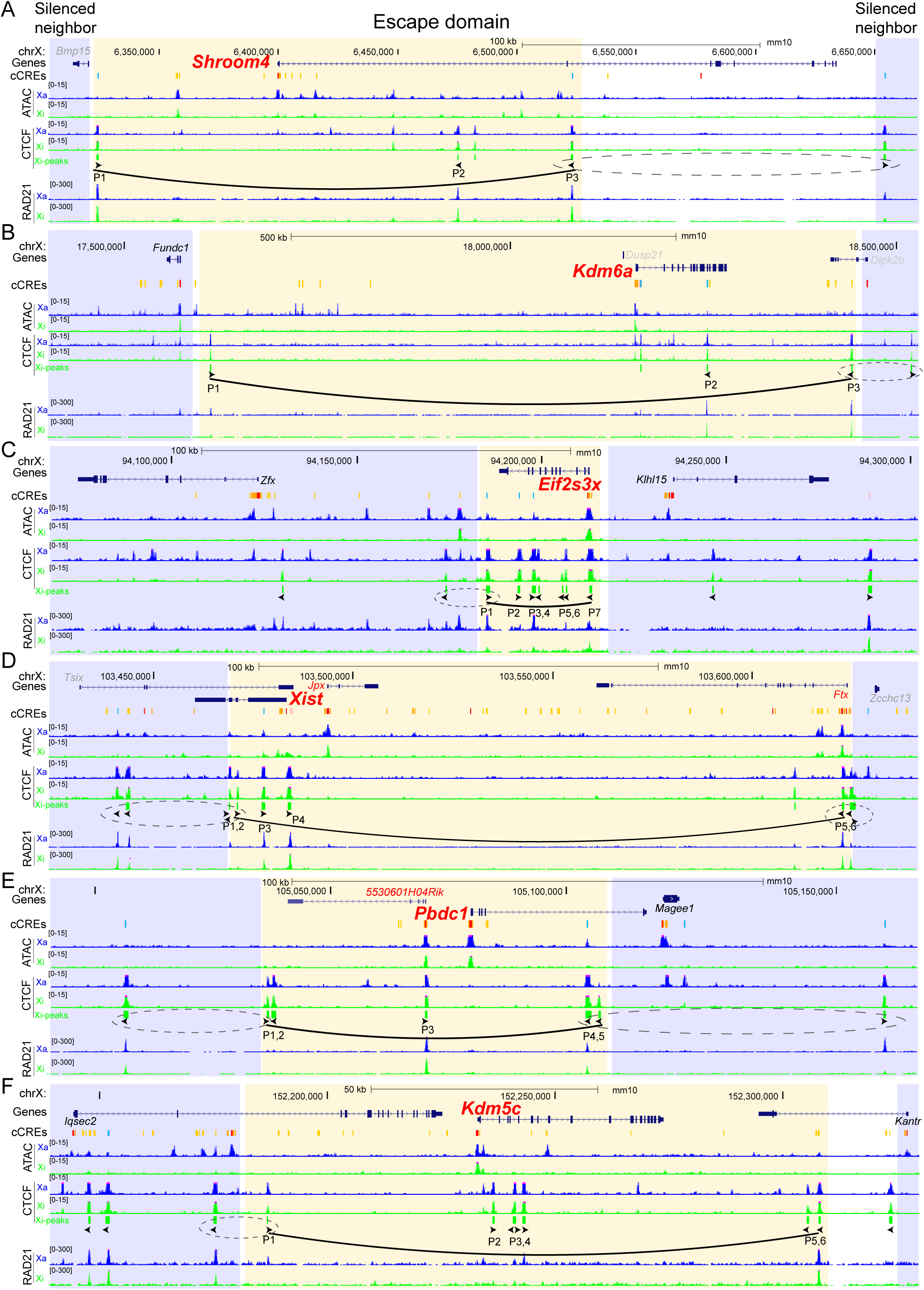
Escape domains in Patski cells. UCSC browser views of domains around the facultative escape gene *Shroom4* (**A**) and the constitutive escape genes *Kdm6a*, *Eif2s3x, Xist, Pbdc1*, and *Kdm5c* (**B-F**). Allelic profiles of ATAC-seq reads and CTCF and RAD21 ChIP-seq reads on the Xa (blue) and Xi (green) are shown. The boundaries between the putative escape domain (yellow) and its flanking silent regions (blue) are marked by several CTCF binding sites with divergent orientation (circled black arrowheads). Inside of the escape domain convergent arrays of CTCF binding sites and RAD21 peaks suggest loop interactions (curve line) mediated by forward motifs and reverse motifs (black arrowheads). ENCODE candidate *cis*-regulatory elements (cCREs) combined for all mouse cell types are shown for promoters in red, enhancers in yellow, and conserved CTCF sites in cyan. Genes known to escape XCI are marked in red, genes subject to XCI, in black, and genes that are not expressed or not assessable due to lack of SNPs, in grey. See Additional file1: Fig. Si for profiles in mouse brain, and Fig. 1A, B for the profiles at the constitutive escape gene *Ddx3x* and facultative escape gene *Car5b* in Patski cells.

It has been reported that CTCF binding site strength is best described by both motif scores and ChIP-seq signals [16]. Thus, we selected CTCF Xi-peaks that have a motif score of at least 10 using CTCFBSDB2.0 [17] and marked their forward or reverse orientations (black arrowheads in Fig. 1A, B, Fig, 2A-F, Additional file 2: Table S2). We found that all eight escapees examined in Patski cells are flanked by convergent arrays of CTCF binding sites (Fig. 1A, B, Fig. 2A-F). The convergent CTCF binding sites may serve as strong anchors of chromatin loops mediated by cohesin to establish escape domains, defined here as putative insulated domains limited by the first forward CTCF binding site at the boundary proximal to the promoter and the last reverse CTCF binding site at the distal boundary. In support of this, RAD21, a cohesin core subunit, often shows peaks that overlap with CTCF peaks at or near the boundaries of escape domains on the Xi (Fig. 1A, B, Fig. 2A-F). Strong and divergent CTCF binding sites were located at 10 of the 16 boundaries between escape genes and adjacent genes subject to XCI (Fig. 1A, B, Fig. 2A-F). In addition, 9/16 boundaries contain one or more CTCF peaks overlapping with cell-type conserved ENCODE CTCF binding sites in mouse [18].

Interestingly, the CTCF binding sites that flank the eight escapees are present on both the Xa and Xi (Fig. 1A, B, Fig. 2A-F). Indeed, metafile analysis of allelic CTCF ChIP-seq signals shows a similar CTCF-binding pattern between the Xa and Xi in Patski cells (Fig. 3A). Thus, the escape domains represent a conserved chromatin structure mediated by the spatial CTCF pattern [16]. Consistent with previous studies [8,11], this indicates that CTCF binding is retained on the Xi during XCI to insulate escape genes from the heterochromatic environment of the Xi. While the putative insulated domains containing the constitutive escapees *Ddx3x*, *Kdm6a*, *Eif2s3x*, *Xist*, *Pbdc1*, and *Kdm5c* are maintained on the Xi both in Patski cells and in adult mouse brain, the domains containing facultative escapees such as *Car5b* and *Shroom4* are only present on the Xi in Patski cells where the genes escape XCI (Fig. 1A, B, Fig. 2A-F, Additional file 1: Fig. S1A). Five of the eight escape domains contain a single escapee, but large escape domains can contain additional escape genes. For example, the *Ddx3x* domain contains *Gpr34* (G-protein coupled receptor 34) that is not expressed in Patski cells, thus precluding defining its escape status in this cell type. However, allelic RNA-seq analysis in mouse brain shows a low level of *Gpr34* escape (Additional file 1: Fig. S1B), which confirms a previous study demonstrating that *Gpr34* is expressed in microglia and escapes XCI in frontal cortex [19]. This illustrates the potential for gene activation within an escape domain when tissue-specific TFs are available. The *Xist* domain (150kb) contains two other escapees, *Jpx* and *Ftx*, both implicated in the process of XCI, while the *Pbdc1* domain contains the lncRNA *5530601H04Rik*, also known to escape XCI (Fig. 2D, E) [20–22].

**Figure 3.**
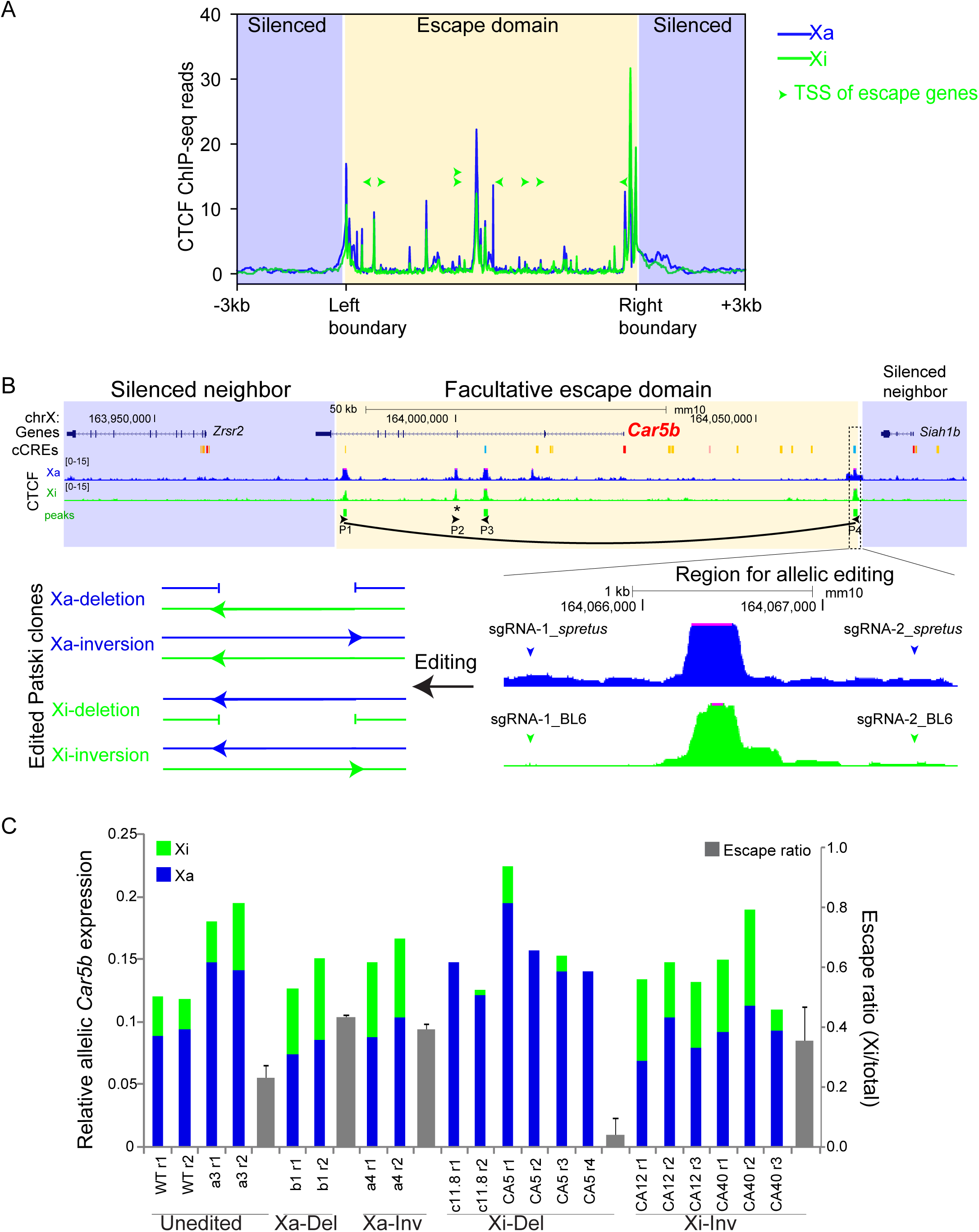
The CTCF boundary region P4 between *Car5b* and *Siah1b* is necessary for *Car5b* escape. **A.** Metafile analysis of CTCF allelic ChIP-seq reads at eight escape domains in Patski cells shows a similar CTCF-binding pattern between the Xa (blue) and Xi (green) (see also Fig. 1A-B, Fig. 2 A-F). The TSS (transcriptional start sites) of escape genes, which are located between CTCF peaks are marked by arrowhead that indicate the direction of transcription. **B**. Schematic of allelic CRISPR/cas9 editing of the 2kb CTCF peak region at P4 to derive deletions and inversions either on the Xi or Xa. Arrows indicate the orientation of CTCF binding sites at P4. A pair of BL6-specific or spretus-specific single-guide RNAs (sgRNAs) was used for editing (Additional file1: Fig. S4, Tables S3 and S4). **C**. Allelic *Car5b* expression from the Xi (green bar) and the Xa (blue bar) measured by quantitative PCR performed on pre-amplified RT-PCR products with or without (mock) ApaI digestion, which specifically cleaves the BL6 allele on the Xi. Relative Xi- and Xa-expression levels are shown in unedited controls (WT and cloned Patski cells in which editing failed), one clone with deletion on the Xa (Xa-Del), one clone with inversion on the Xa (Xa-Inv), two independent clones with deletion on the Xi (Xi-Del), and two independent clones with inversion on the Xi (Xi-Inv). At least two biological replicates were tested for WT Patski cells and for each cloned line. Escape levels of *Car5b* (grey bar) calculated from the Xi/total expression ratios are shown as means ± SEM. Only the Del-Xi lines show a significant decrease in escape levels, as compared to any other group (P < 0.0005, unpaired two-tail *t*-test). Note that Xi-specific expression ranges between 20-40% of total *Car5b* expression.

We conclude that the majority of escape domains are flanked by convergent CTCF binding sites present on both the Xa and Xi. For constitutive escape genes this configuration remains unchanged in Patski cells and adult mouse brain, but for facultative escape genes CTCF sites are only present in cells where the gene escapes XCI.

### Characterization of escape domains

We next examined the relationships between the escape domains and topologically associating domains (TADs). We found that the putative insulated domains limited by convergent CTCF binding sites around escapees are usually smaller than the TADs we previously defined on the Xi using allelic contacts at 520kb windows with 40kb resolution (Fig. 1A, B) [15]. Escape domains, whose size is usually less than 200kb, are often located inside TADs and thus could represent allelic sub-TAD structures or loops (Additional file 2: Table S1). Two exceptions are the *Ddx3x* domain that measures ∼500kb and overlaps with a known TAD in both Patski and brain (Fig. 1A, B) [15] and the *Kdm6a* domain that measures ∼800kb and overlaps a very large TAD in Patski cells and a smaller one in brain (Additional file 2: Table S1).

The spatial distribution of CTCF binding sites was then mapped relative to the location of the promoter and enhancers of each of the eight escapees within the putative insulated domains. For four of the escapees, *Shroom4*, *Xist*, *Pbdc1*, and *Car5b,* CTCF arrays flank the promoter and enhancers but not the 3’ end. Rather, the convergent CTCF arrays are located within each of these genes, suggesting the formation of loops that mainly insulate the promoter region. Accordingly, the TSS (transcriptional start site, i.e., the center of the promoter; green arrowhead in Fig. 3A) of each escapee is often located between CTCF peaks, which would allow access of transcription factors (TFs), interaction with enhancers, and gene activation in a critical region within escape domains.

Allelic ATAC-seq and CUT&RUN analyses were done to map chromatin accessibility and histone modifications at escape domains. Except for *Xist*, which is only expressed from the Xi, we found that chromatin accessibility is reduced at all eight escape domains on the Xi compared to the Xa (Fig 1A, B, Fig. 2A-F). Enrichment in the active enhancer mark H3K27ac is similarly reduced (Additional file 1: Fig. S2), consistent with reduced levels of RNA polymerase and lower expression of escapees from the Xi versus the Xa [8,10,23,24]. As expected, the repressive histone mark H3K27me3 is depleted at all eight escape genes on the Xi (Additional file 1: Fig. S2). Interestingly, the strong CTCF peaks at the boundary of escape domains often have no or weak ATAC-seq peaks on both the Xi and the Xa (Fig. 1A, B, Fig. 2A-F), suggesting a tight chromatin structure protected by strong CTCF binding. Conversely, the strong ATAC-seq peaks at the promoters of escape genes (e.g., *Pbdc1*, *Kdm5c*) often have no or weak CTCF peaks, consistent with the TSS of escapees being often located between CTCF peaks (Fig. 3A). This is in line with previous findings that YY1, not CTCF, is often a major regulator of promoter-enhancer interactions [25].

In summary, CTCF sites are often located proximal to the promoter and within escape genes, suggesting insulation of the promoter region. Chromatin accessibility and active histone marks are associated with expression of escapees on the Xi, albeit at a reduced level compared to the Xa.

### *Car5b* is protected from XCI by CTCF insulation

To test the function of CTCF in regulating escape from XCI, we focused on the tissue-specific escape gene *Car5b* (Fig. 1B). *Car5b/CA5B* encodes a highly conserved mitochondrial carbonic anhydrase expressed in multiple human and mouse tissues with increased expression during development and high expression in kidney (Additional file 1: Fig. S3A-B). In human *CA5B* escapes from XCI in most tissues [6]. In contrast, *Car5b* is subject to XCI in mouse adult tissues including brain, spleen, heart, kidney, liver and ovary as well as in MEFs (Additional file 1: Fig. S3C-D). However, *Car5b* escapes XCI in Patski cells that were derived from E18.5 embryonic kidney, which provides a cell line system for editing the escape domain.

CTCF binds to a region proximal to the *Car5b* promoter on the Xi in Patski cells but not in brain, suggesting that CTCF-mediated insulation protects the *Car5b* escape domain in Patski cells (Fig. 1B) [8]. This CTCF binding peak (hereafter P4; marked in Fig. 1B) is located at the boundary between *Car5b* and the neighboring gene *Siah1b* that is subject to XCI. This is a conserved CTCF binding region as shown by *cis*-element predictions from ENCODE [18], which includes a binding site with a high reverse motif score of 26 (chrX:164066195-164066766; Additional file 2: Table S2). We hypothesized that *Car5b* escape may be due to the formation of a loop domain, supported by our findings of RAD21 binding and by virtual 4C analysis based on Xi-specific Hi-C contact maps that show interactions between P4 and three CTCF Xi-peaks (P1-3) located within *Car5b* (Fig. 1B). To test the role of P4 we deleted or inverted it in Patski cells using allele-specific sgRNAs for CRISPR/Cas9 editing (Fig. 3B, Additional file 1: Fig. S4A, Additional File 2: Table S3). At least two independent clones were derived for each type of heterozygous editing on the Xi (Xi-Del and Xi-Inv) and on the Xa (Xa-Del and Xa-Inv). Correct editing of the targeted alleles and retention of the intact CTCF site on the unedited alleles were confirmed by PCR and Sanger sequencing (Additional file 1: Fig. S4B-C). The intact alleles in each clone were used as internal controls for gene expression and epigenetic analyses.

Using allelic expression analysis based on BL6-specific ApaI digestion [21] we determined that *Car5b* escape was abolished or strongly reduced following deletion of P4 in Xi-Del clones, while inversion of this same region in Xi-Inv clones had no effect (Fig. 3C, Additional file 1: Fig. S5A-B). No reactivation of the neighbor gene *Siah1b* was observed (Additional file 1: Fig. S5C-D). In contrast, no change in *Car5b* expression was observed when P4 was deleted or inverted on the Xa (Fig. 3C, Additional file 1: Fig. S5A-B). This indicates that CTCF binding at the boundary region P4 is essential for escape from XCI on the Xi but not for gene expression on the Xa.

Next, we generated allelic profiles for CTCF, H3K27ac, and H3K27me3 by CUT&RUN in Xi-Del, Xi-Inv, and Xa-Del lines, and for RAD21 by ChIP-seq in wild-type (WT) and Xi-Del lines. Deletion of the CTCF region P4 on the Xi results in reduced levels of CTCF and H3K27ac and in increased levels of H3K27me3 over the *Car5b* escape domain on the Xi, consistent with silencing (Fig. 4A-D, Additional file 1: Fig. S6A-C). In contrast, no apparent changes are present when P4 is inverted on the Xi or deleted on the Xa (Fig. 4A-C). Changes on the Xi are more pronounced in clone Xi-Del-c11.8, where there is complete loss of escape, than in clone Xi-Del-CA5, where a variable and very low level of escape persists (Fig. 3C, Fig. 4A-D, Additional file 1: Fig. S5B, S6A-C). In clone Xi-Del-c11.8 loss of active marks (H3K27ac) and spreading of silencing marks (H3K27me3) is evident throughout the whole escape domain (∼85kb) (Fig. 4B, C).

**Figure 4.**
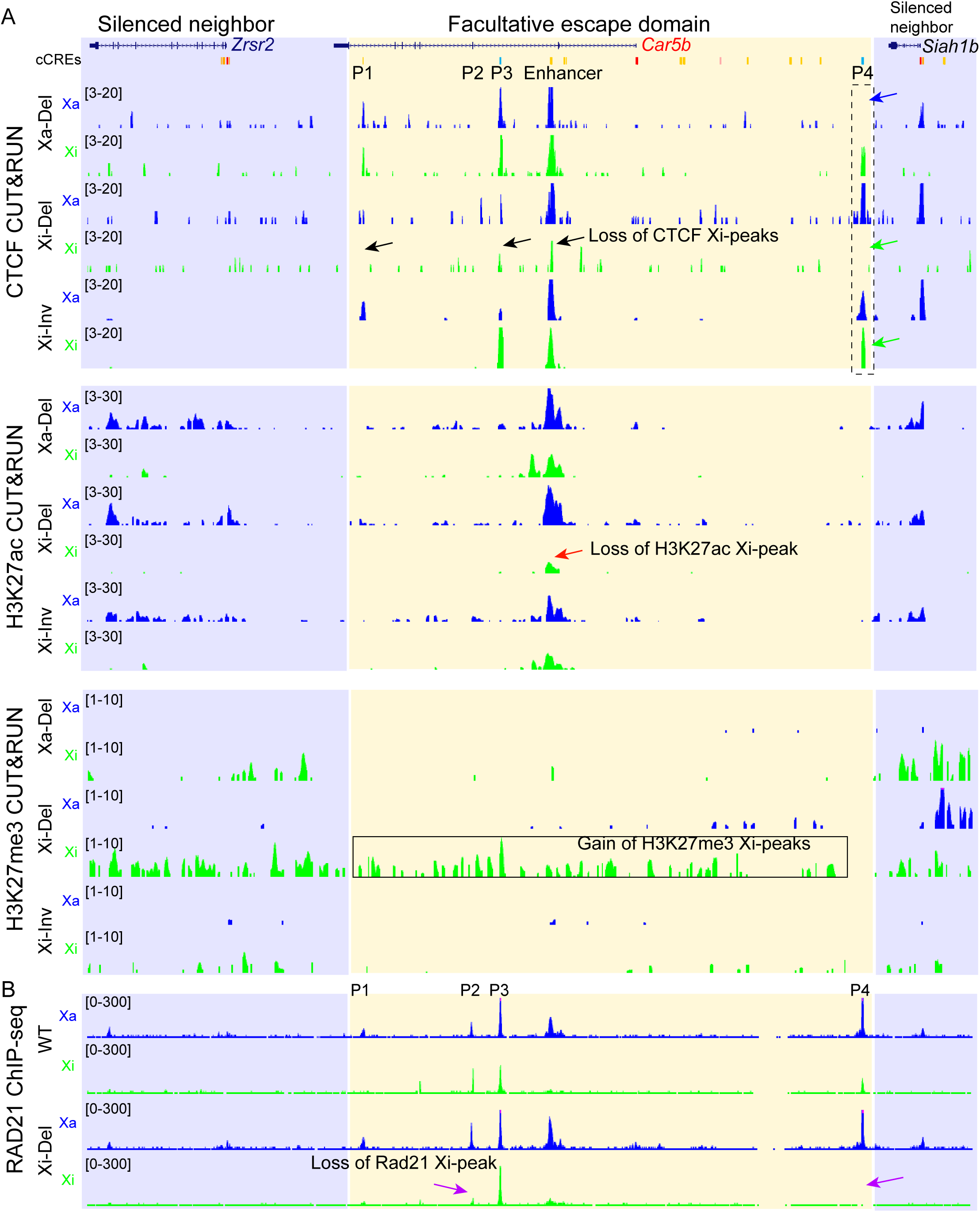
CTCF binding protects *Car5b* from heterochromatic marks on the Xi. **A.** UCSC browser view of allelic profiles of CUT&RUN reads for CTCF, H3K27ac, and H3K27me3 at the *Car5b* escape domain (yellow) and surrounding silent regions (blue) in Xa-Del (clone b1), Xi-Del (clone c11.8) and Xi-Inv (clone CA12). The expected loss of CTCF binding due to editing is seen at the P4 region (blue and green arrows). In Xi-Del the CTCF binding sites P1 and P3 (black arrows) and the active enhancer mark H3K27ac (red arrow) are largely reduced on the Xi. Importantly, the silencing mark H3K27me3 (gray box) is gained throughout the *Car5b* domain, suggesting that CTCF functions as an insulator to protect escape. No apparent changes were observed at *Car5b* in Xa-Del or Xi-Inv cells. **B.** UCSC browser view of allelic RAD21 ChIP-seq reads in WT and Xi-Del Patski cells. There is a loss of RAD21 peaks at the edited P4 region and also a decrease at the P2 region (marked by asterisk in Fig. 1B) inside *Car5b* (purple arrows) on the Xi in Xi-Del. No changes were observed on the Xa.

Two (P1, P3) of the three sites with reduced CTCF binding within the *Car5b* locus after deletion of P4 overlap with chromatin contact regions identified by virtual 4C (Fig. 1B, Fig. 4A). The third site (P2) with reduced CTCF binding overlaps with an enhancer region identified by ATAC-seq and H3K27ac peaks and confirmed as an ENCODE annotated *cis*-regulatory enhancer (Fig. 1B, Fig. 4A). In clone Xi-Del-CA5 only P1 and the enhancer showed reduced CTCF binding (Additional File 1: Fig. S6A). Note that CTCF binding at the enhancer is captured by CUT&RUN that can detect both bound regions and regions located in proximity of a binding site [26]. However, ChIP-seq, which can only detect bound regions, did not reveal a signal at the enhancer, suggesting that the enhancer is not directly bound by CTCF but is located near a CTCF binding region (Fig. 1B, Fig. 4A, Additional File 1: Fig. S6A). In addition to the expected loss of RAD21 binding at deleted P4 in clone Xi-Del-c11.8 we also observed loss of RAD21 at site P2 that contains a forward CTCF motif and thus could mediate a smaller loop (Fig. 1B, Fig. 4D).

Our allelic analysis of gene expression and epigenetic features in cell lines with allele-specific deletions or inversions of a CTCF boundary strongly support the role of CTCF in protecting the *Car5b* escape domain on the Xi via boundary and looping functions.

### Escape genes are sensitive to disruptions of the Xi-specific heterochromatic environment

We previously reported that either reducing Xi-specific H3K27me3 enrichment by depletion of the *Firre* lncRNA (Del-Firre; [27]) or disrupting the Xi compact structure by deletion of the hinge containing the *Dxz4* locus (Del-Hinge; [15]) results in increased expression of a small set of genes from the Xi. In particular, *Car5b* expression is increased 2.3-fold and 1.6-fold by *Firre* RNA depletion and hinge deletion, respectively (Fig. 5A-B). Interestingly, ectopic expression of a *Firre* cDNA transgene completely restores normal *Car5b* escape levels, consistent with the reversible trans-acting effect of *Firre* RNA on H3K27me3 enrichment on the Xi (Fig. 5A) [27]. At other escapees we also found increases in Xi-expression levels, which are more pronounced after depletion of *Firre* RNA than after deletion of the hinge (Fig. 5C). These results indicate that the heterochromatic environment of the Xi, including enrichment in H3K27me3 and chromatin condensation, normally constrain the level of expression from escapees. Since the escapees are distributed along the whole Xi, these effects appear to be global.

**Figure 5.**
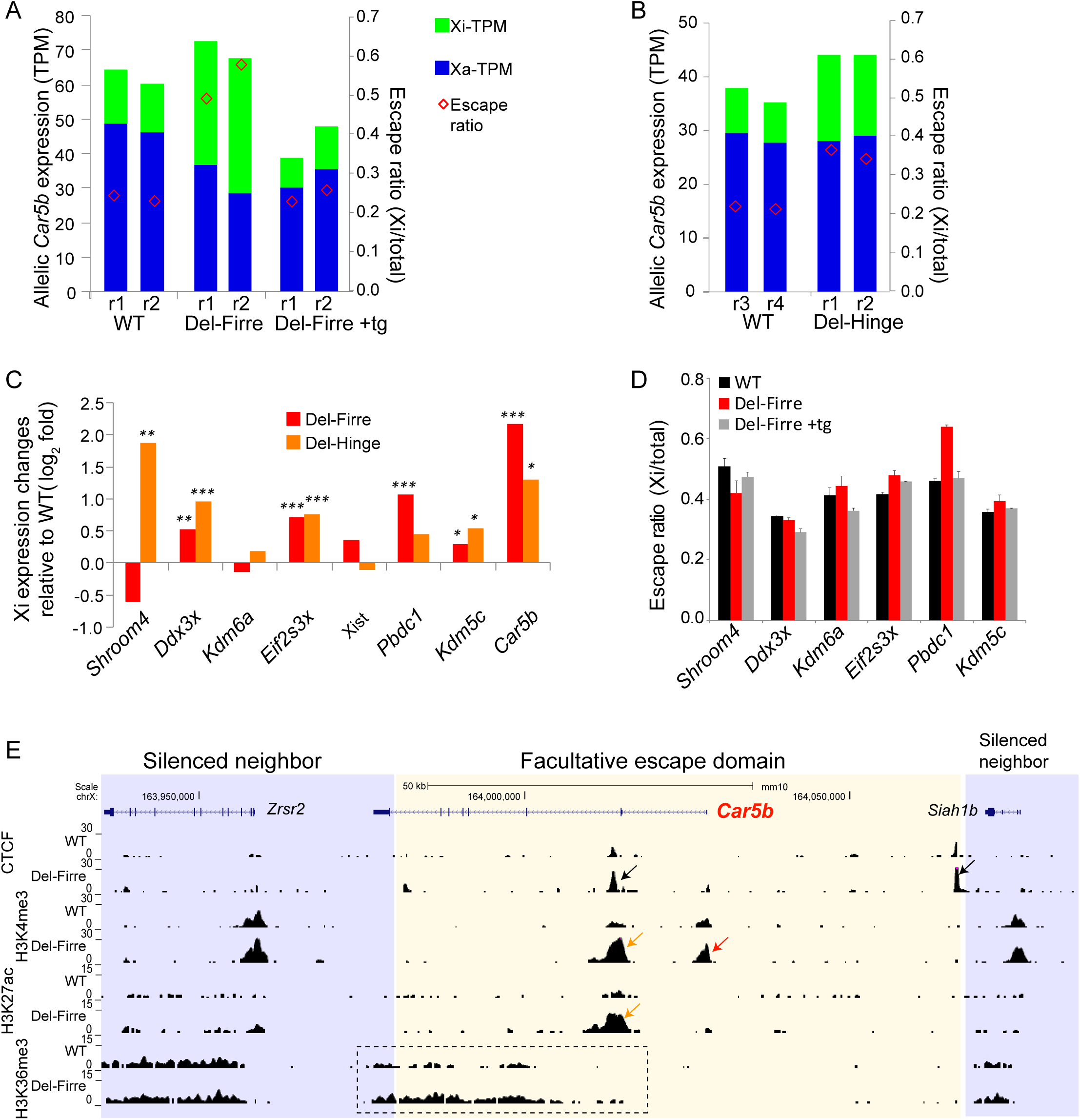
The chromatin environment constrains expression levels of escape genes on the Xi. **A**. Depletion of *Firre* RNA in Patski cells by deletion of the lncRNA on the active X chromosome (Del-Firre), which causes a loss of H3K27me3 on the Xi, increases escape levels of *Car5b* that can be reversed using a *Firre* cDNA transgene (Del-Firre+tg). Allelic expression levels (TPM, transcript per million) for the Xa (blue) and Xi (green) are shown together with escape ratios (red diamond). RNA-seq data from Fang et al., 2020 [27]. **B**. Disruption of the Xi condensed structure by deletion of the *Dxz4* locus on the Xi (Del-hinge) also increases *Car5b* expression on the Xi. Same analysis as in A. RNA-seq data from Bonora et al., 2018 [15]. **C.** Both depletion of *Firre* RNA and deletion of the *Dxz4* locus increase Xi-expression levels of eight escape genes as shown by Xi-expression fold changes compared to WT controls. Fold changes and adjusted P-values were derived from allelic DESeq2 analysis of RNA-seq data. **D.** Except for *Shroom4*, increased escape levels of five escape genes upon *Firre* RNA depletion (Del-Firre) are restored by ectopic expression of a *Firre* cDNA transgene (Del-Firre+tg). **E.** UCSC browser view of total reads from CTCF, H3K4me3, H3K27ac, and H3K36me3 CUT&RUN in WT and Del-Firre Patski cells. Note that allelic analysis was not possible due to limited coverage and short reads (36nt) in this CUT&RUN batch. Considering that expression of *Car5b* on the Xi but not on the Xa is significantly increased in Del-Firre cells as shown by allelic RNA-seq in A, we infer that increases in CTCF (black arrows) and in active histone marks at the promoter (H3K4me3; red arrow), the enhancer (H3K4me3 and H3K27ac; orange arrows), and 3’ region (H3K36me3 for elongation; dotted box) of *Car5b* is probably from the Xi allele.

Consistent with increased *Car5b* Xi-expression due to loss of *Firre* RNA, Xi-specific profiles of active marks by CUT&RUN show an increase in H3K4me3, H3K27ac, and H3K36me3 (Fig. 5E). Note that due to limited SNP read coverage in this dataset allelic analyses were not feasible (See Methods). However, since *Car5b* expression on the Xa was not changed after loss of *Firre* RNA, we speculate that the observed increase in active marks occurs on the Xi allele (Fig. 5A, C-E). Cells with deletion of the Xi-specific hinge showed a moderate increase in chromatin accessibility at *Car5b* on the Xi as measured by ATAC-seq (Additional file 1: Fig. S7A), consistent with a modest expression increase (Fig. 5B, C). Since chromatin organization of the Xi becomes more like that of the Xa in Del-Hinge cells [15], we also edited the same CTCF boundary region P4 in Del-Hinge cells to test whether the boundary is still required for *Car5b* escape. Indeed, deletion of P4 on the Xi in Del-hinge cells strongly reduces *Car5b* escape, similar to what is observed in Xi-Del cells derived from WT (Additional file 1: Fig. S7B). Thus, the CTCF boundary is required for *Car5b* escape whether the hinge is intact or disrupted, consistent with the notion that *Xist* coating and the silencing mark H3K27me3 are not affected by Del-hinge [15]. The increase in *Car5b* expression from the Xi in Del-Hinge would then be due to lower chromatin compaction of the Xi.

Next, we tested whether inhibition of DNA methylation that is essential for silencing genes subject to XCI has any effect on *Car5b* escape levels by examining our previous published RNA-seq data [15]. Inhibition of DNA methylation using 5-aza-cytidine had no effect on *Car5b* escape levels and only caused minor reactivation of neighbor genes *Zrsr2* and *Siah1b* that are subject to XCI in both WT and Del-hinge cells (Additional file 1: Fig. S8A-C). Interestingly, reactivation levels of these neighbor genes appear to be positively related to their distance from the boundaries of the escape domain (closer to *Siah1b* than *Zrsr2)*. In addition, escape levels of the neighbor genes were much lower compared to that of *Car5b*, consistent with limited reactivation, which was also shown in a previous report of only ∼2% of cells with reactivation of a reporter after inhibition of DNA methylation [28].

These findings indicate that the 3D structure and repressive heterochromatic marks of the Xi but not DNA methylation play a role in modulating expression levels of escapees on the Xi.

## Discussion

CTCF has been proposed since 2005 to play an important role in regulation of genes that escape XCI [11]. Insulation via CTCF binding would partially protect escape genes from the Xi silencing environment and thus allows access to TFs for gene transcription, albeit at a reduced level compared to the Xa allele. While this hypothesis is supported by a few transgenic studies [12–14], our study is the first to report that convergent arrays of CTCF binding sites insulate an escape gene, most likely by stabilizing cohesin-mediated chromatin loops. We find that CTCF binding around the escape gene *Car5b* is not Xi-specific since it also exists on the Xa, suggesting these CTCF binding sites may be specifically retained at escape genes on the Xi during the establishment of XCI. This retention process or possibly loss followed by re-acquisition may be tissue-specific, which would result in genes with facultative escape. While CTCF clearly plays an important role in establishing chromatin structure during embryogenesis [29], future studies are needed to track allelic CTCF binding profiles during development, especially at facultative escape genes to understand how CTCF binding is retained or lost.

Our results show complete loss of *Car5b* escape when the CTCF boundary region P4 is deleted on the Xi in Patski cells, while in brain where the gene does not escape XCI there is no evidence of CTCF binding around *Car5b*. In contrast, expression of *Car5b* from the Xa where chromatin is opened does not depend on CTCF binding, as shown by deletion of region P4 on the Xa. This is consistent with previous findings that show no or limited (often locus-specific) effects on gene expression after either deletion of a CTCF site that disrupts a chromatin loop or acute depletion of CTCF protein [30,31]. Thus, CTCF may be mainly involved in the insulation of chromatin structures and play a limited role in directly regulating gene expression. In contrast, YY1 plays a direct role in regulation of promoter-enhancer interactions [25] and is enriched at the TSS of escapees [32]. Our findings of no or weak CTCF binding at promoters and enhancers of escape genes confirm these reports. This is also consistent with the recent finding that there is no significant difference in predicted probability of CTCF binding at the TSS of escape genes versus genes subject to XCI in human and mouse [7]. Interestingly, we found that *Car5b* escape depends on CTCF binding region P4 proximal to the promoter region, which interacts with CTCF sites P1-3 within the gene. Thus, the insulated domain includes the promoter and enhancer of *Car5b* but lacks its 3’ end. This is also observed at *Shroom4*, *Xist*, and *Pbdc1.* Thus CTCF-mediated chromatin boundaries do not stall the machinery for transcription elongation [16].

Based on peak analysis, CTCF binding sites appear to be stronger at the boundary between escape genes and neighbor genes subject to XCI compared to those within escape domains. Furthermore, these boundary sites often correspond to cell-type conserved CTCF regulatory sites and overlap with RAD21 binding, suggesting that they represent master *cis*-elements to anchor cohesin loops for domain formation. Our results show that deletion of the CTCF boundary region P4 causes several changes in histone modifications associated with changes in gene expression. In addition, there is no spreading of escape into the neighbor gene *Siah1b* upon deletion of the CTCF boundary at *Car5b* on the Xi. This is different from results of a transgenic study showing that insertion of the escape gene *Kdm5c* without its right boundary located inside the neighbor gene *Kantr* (see Additional file: Fig. S1F) causes ectopic escape of genes downstream of the insertion [12]. Whether this difference is due to different effects from deletion of the endogenous *Car5b* boundary site versus deletion of a BAC transgene is unclear. Alternatively, regulatory elements associated with constitutive escape genes (*Kdm5c*) could be different from those of compared facultative escape genes (*Car5b*), leading to different effects upon deletion of boundaries. Further editing of boundaries of other escape domains is needed to examine this question. Intriguingly, we found that inversion of region P4 on the Xi had no effect on *Car5b* escape, suggesting that coiled looping, which has been reported to occur genome-wide [33], could indeed be mediated via tandem CTCF binding sites and still function as insulation. The strength of loop anchoring via a pair of tandem CTCF binding sites appears to be lesser than via a pair of convergent CTCF binding sites. However, a recent study has shown that even without consideration of CTCF binding motif orientation a pair of dCas-fused CTCF molecules can induce insulation and enhance escape from XCI [14].

We have previously mapped the position of escapees at the periphery of the 3D Xi-structure in regions enriched in CTCF binding [34], which is consistent with our current findings of putative insulated domains of escapees on the Xi. Indeed, escapees have been reported to show increased inter-chromosomal contracts compared to genes subject to XCI [35]. However, our findings of reduced levels of chromatin accessibility and active marks at escape domains on the Xi versus the Xa and of an increase in escape levels in cells with disrupted H3K27me3 or de-condensation of the Xi suggest that escape domains are under spatial constraints that limit access to TFs, resulting in lower expression levels from the Xi versus the Xa. Escape levels are probably controlled by multiple regulatory layers, including *Firre*-mediated H3K27me3 maintenance [27] and *Dxz4*-mediated 3D structure [15]. In addition, escape levels in somatic cells are known to be sensitive to *Xist* RNA levels [36,37]. Interestingly, escapees may be sensitive to different types of regulation. For example, we found that loss of H3K27me3 in a *Firre* mutant resulted in full escape of *Car5b* and *Pbdc1,* but not for other escapees whose expression from the Xi remained lower than on the Xa. We also note that loss of H3K27me3 affected escape levels to a greater extent than Xi de-condensation. Escape regulatory mechanisms could also differ depending on whether alterations are made before or after the onset of XCI. For example, deletion of *Dxz4* in stem cells prior to the onset of XCI causes loss of escape of the facultative escapee *Mecp2* once the cells are differentiated [38], whereas our findings in Patski cells in which XCI is established increases levels of escape. It will be interesting to follow the dynamic, regulatory interactions of escape during development.

## Conclusion

Our findings strongly support the role of insulation and looping via convergent arrays of CTCF binding sites in maintenance of escape from XCI. Our study shows that escape from XCI is further modulated by the 3D structure of the Xi and its H3K27me3 enrichment but not by DNA methylation. These findings provide insights for future studies that aim to perturb gene silencing/escape on the Xi to help understand their roles in human health and diseases.

## Methods

### Mouse tissues and cell lines

Female F1 hybrid progeny were obtained by mating C57B/6J females that carry a deletion of the *Xist* proximal A-repeat (*Xist^ΔA^*) (B6.Cg-Xist, RIKEN) with *Mus spretus* males (Jackson Labs) as described [8]. Mouse adult tissues and embryonic fibroblasts (MEFs) were collected from F1 mice or embryos that inherited a maternal X chromosome with an *Xist^ΔA^*. These *Xist^ΔA^*/+ tissues and MEFs fail to silence the BL6 X and thus have complete skewing of XCI of the paternal *spretus* X. All procedures involving animals were reviewed and approved by the University Institutional Animal Care and Use Committee (IACUC), and were performed in accordance with the Guiding Principles for the Care and Use of Laboratory Animals. Patski cells are fibroblasts previously derived from 18dpc embryonic kidney from a cross between a BL6 female mouse with an *Hprt^BM3^* mutation and *M. spretus* male [39]. The cells were selected in HAT media such that the BL6 X chromosome is always inactive as verified in previous studies [8,23]. Patski cells were cultured as previously described [23].

### Allele-specific CRISPR/Cas9 editing of Patski cells

Allele-specific CRISPR/Cas9 editing of the CTCF binding region upstream of *Car5b* was performed in Patski cells as described previously [15]. Allele-specific sgRNAs designed using CHOPCHOP were selected to include BL6 or *spretus* SNPs at multiple sites or at the PAM site depending on availability (Additional file 1:Fig. S4A, Additional file 2: Table S3). Patski cells were transfected using Ultracruz transfection reagents (Santa Cruz). PCR together with Sanger sequencing was used to verify specific deletion or inversion of the BL6 or *spretus* allele in each clone and to verify junction sequences containing BL6 SNPs (Additional file 1: Fig S4B). WT Patski cells and clones that had been subjected to transfection but did not carry any deletions or inversions were used as unedited controls. The sequence of the primers used is in Additional file 2: Table S4.

### Allelic expression analysis using BL6-specific restriction enzyme digestion

Allelic expression analysis of *Car5b* was done by BL6-specific ApaI digestion as described [23]. In brief, cDNA obtained by reverse transcription (RT) of RNA and genomic DNA were amplified by PCR using primers flanking a BL6-specific ApaI recognition site in exon 4 of *Car5b*. PCR products were then purified followed by mock (-ApaI) or ApaI digestion and gel electrophoresis. In Patski cells the appearance of a digested band indicates expression from the BL6 allele on the Xi and thus escape from XCI (see Additional file 1: Fig. S5B). In contrast, in mouse tissues and MEFs the presence of an undigested band would indicate expression from the *spretus* allele on the Xi and thus escape from XCI (see Additional file 1: Fig. S3C). Allelic expression analysis of *Siah1b* was done similarly using BL6-specific BsrI digestion (see Additional file 1: Fig. S5C-D). The sequence of primers used is in Additional file 2: Table S4.

To quantify escape levels, quantitative PCR was performed using a SYBR green system for pre-amplified RT-PCR products with mock or BL6-specifc ApaI digestion as described in [23]. Only undigested products can be amplified and measured, allowing the measurement of the difference between mock and BL6-specifc ApaI-digested samples as an indicator of escape levels. At least two biological replicates were tested for each cloned lines.

### ATAC-seq, ChIP-seq, CUT&RUN, Hi-C

ATAC-seq, CTCF ChIP-seq, and Hi-C datasets as well as allelic data analyses are described in our previous studies [8,15,27]. Virtual 4C analysis was performed as described [15]. ChIP-seq for RAD21 was performed in WT and CTCF-site edited Patski cells using a rabbit monoclonal antibody for RAD21 (Abcam ab217678) as described previously [8]. CUT&RUN was performed in WT and CTCF-site edited Patski cells using antibodies for CTCF (Millipore 07-729), H3K27ac (Abcam ab4729) and H3K27me3 (Millipore 07-449), as described in [27]. Allelic data analysis was done as described [15]. Additional CUT&RUN datasets of CTCF, H3K4me3, H3K27ac3, and H3K36me3 in WT and *Firre*-deleted Patski cells were obtained from Thakur et al. [40]. Note that no allelic analyses were possible due in this batch to limited SNP coverage in short (36nt) reads.

### CTCF motif analysis

CTCF Xi-peak sequences (100-200bp) in Patski cells were used to scan for conserved CTCF motifs, including detection of the Ren_20 motif and the MIT long motifs (LM2, LM7 and LM23), and determination of motif scores and orientation by CTCFBSDB 2.0 (insulatordb.uthsc.edu, [17]) (Additional file 2: Table S2). Only CTCF Xi-peaks with a motif score ≥10 and clear orientation were considered to evaluate the CTCF binding pattern around escape genes. Note that for peaks containing two or more CTCF binding sites with a motif score ≥10, all motif sites are marked on the figures. Using a motif score >3 cutoff, which is suggestive of a CTCF binding site, we obtained similar results.

## Acknowledgements

We thank lab members for their help in this study and we thank Charlie Lee for his help with next-generation sequencing.

## Author’s contributions

XD and CMD designed the experiments; HF, ART, TN, JT and XD performed the experiments; HF, ART, GB, and XD analyzed the data; JB, GNF, SH, JS, WSN provided materials, data, and/or analysis support; XD and CMD wrote the paper. All the authors read and approved the final manuscript.

## Availability of data and materials

High-throughput sequencing data that support the findings of this study have been deposited in the National Centre for Biotechnology Information GEO and are accessible through the GEO Series “GSE59779” and “GSE231626”. All other data and the scripts used for the analyses that support the findings of this study are available within the article and its Supplementary Information files or from the corresponding authors upon reasonable request.

## Additional file 1

**Figure S1.**
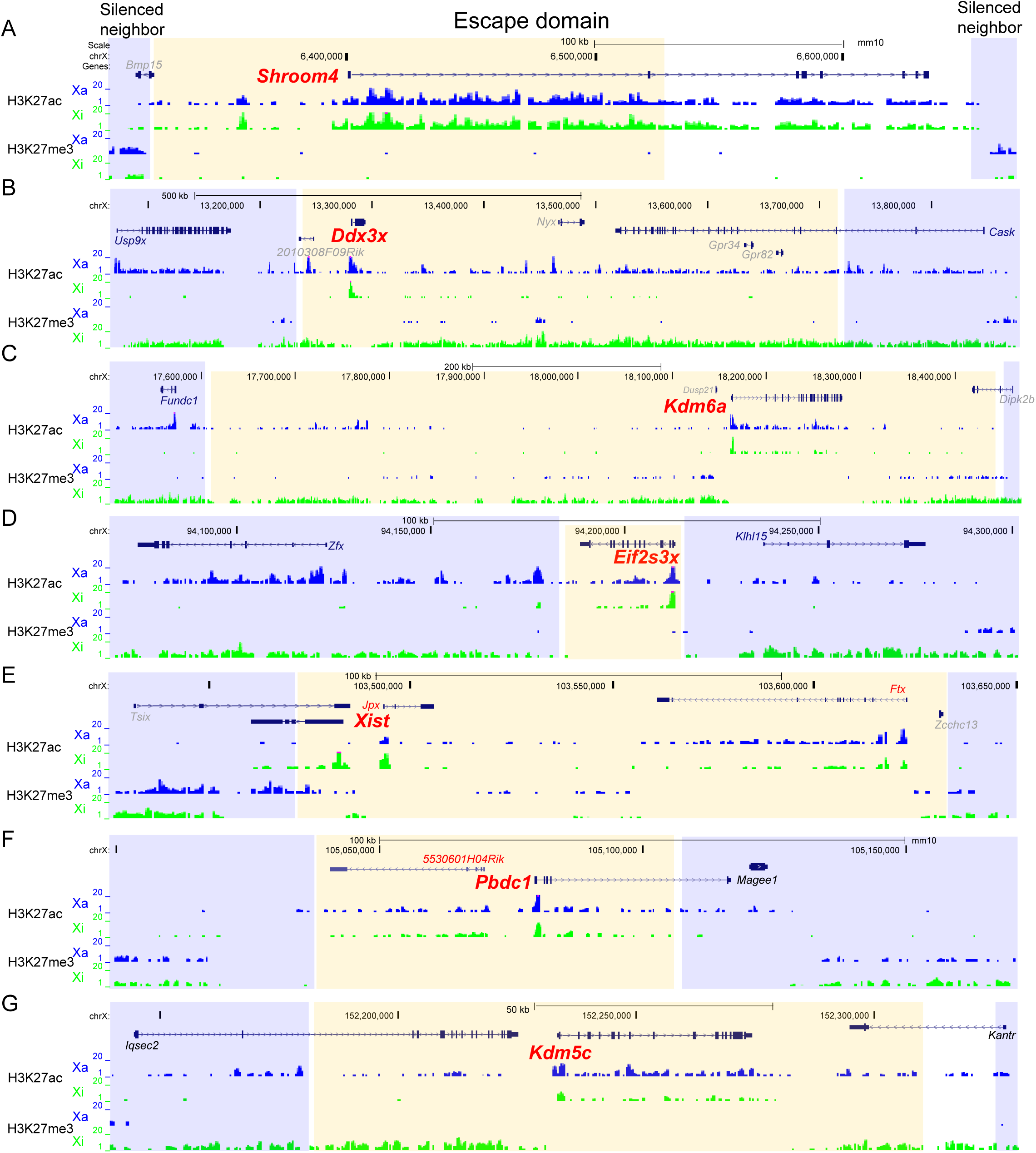
Escape domains in adult mouse brain. **A.** UCSC browser views of profiles of CTCF ChIP-seq reads on the Xa (blue) and Xi (green) are shown in domains around the facultative escape gene *Shroom4* and the constitutive escape genes *Kdm6a*, *Eif2s3x*, *Xist*, *Pbdc1*, and *Kdm5c* in adult mouse brain. Putative escape domains are shaded in yellow, and silenced domains in blue. While the CTCF peaks that flank escape domains around constitutive escape genes as defined for *Ddx3x* in Patski cells (Fig. 1A) are also present in brain, CTCF peaks are absent around the facultative escape gene *Shroom4*, which is silenced in brain, similar to *Car5b* (Fig. 1B). ChIP-seq data from Berletch et al., 2015). Escape genes are in red, inactivated genes in black, and genes with an unknown XCI status, in grey. See also Fig. 2A-F. **B.** Low levels of *Gpr34* expression from the Xi are apparent in mouse adult brain, indicating escape from XCI. UCSC browser view of RNA-seq reads on the Xa (blue) and Xi (green) are shown at *Gpr34*.

**Figure S2.**
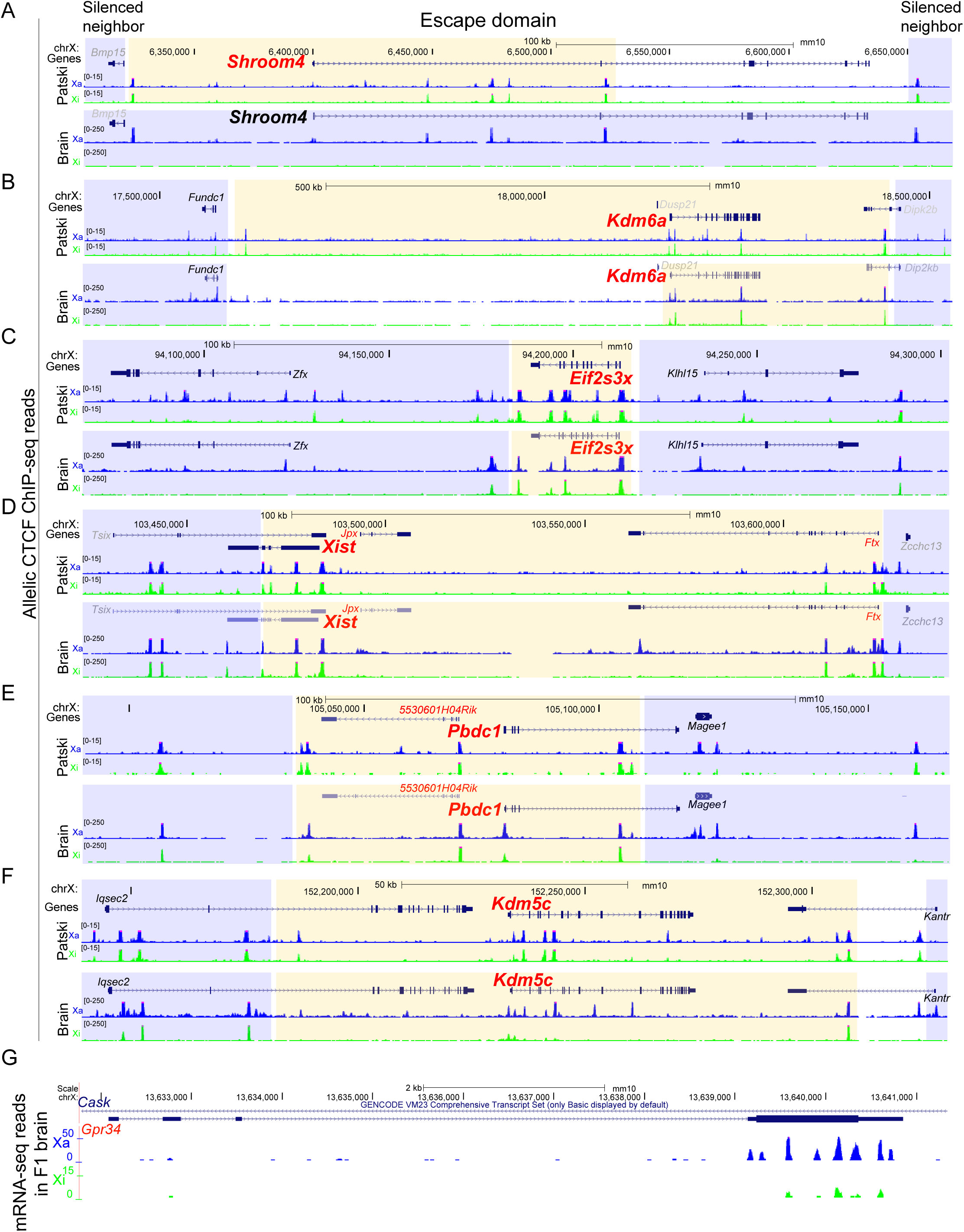
Allelic profiles of histone marks at escape domains in Patski cells. UCSC browser views of profiles of CUT&RUN reads on the Xa (blue) and Xi (green) are shown for H3K27ac as an active enhancer mark and H3K27me3 as an repressive mark at the escape domains of (**A**) the facultative escape gene *Shroom4* and (**B-G**) the six constitutive escape genes, *Kdm6a*, *Ddx3x, Eif2s3x*, *Xist*, *Pbdc1*, and *Kdm5c*. Escape genes show enrichment of H3K27ac and depletion of H3K27me3 on the Xi, while neighbor genes subject to XCI show an opposite pattern. Putative escape domains are shaded in yellow, and silenced domains in blue. See Fig. 4A-C for profiles at *Car5b*.

**Figure S3.**
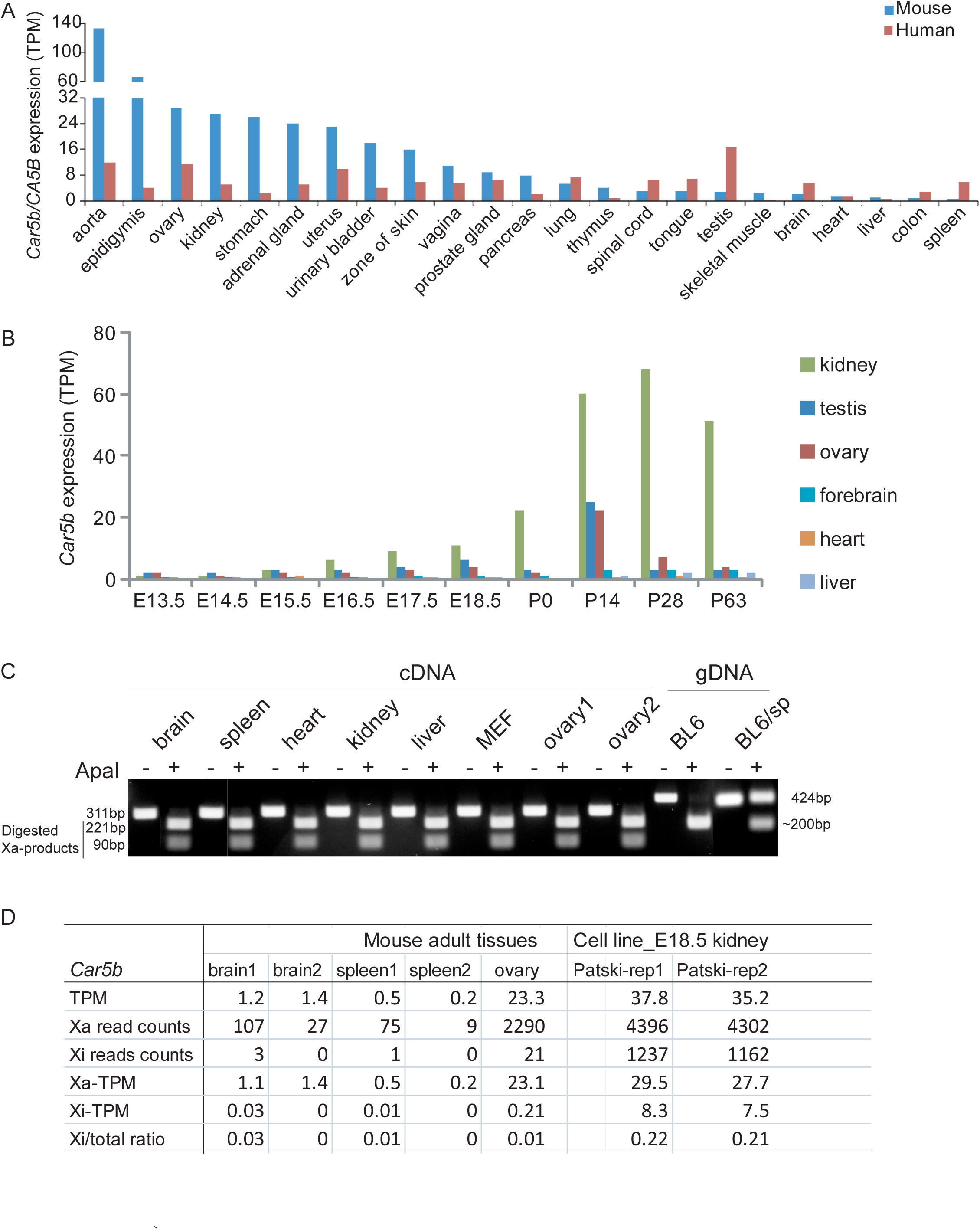
Conserved expression of *Car5b* in mouse and human tissues and escape status. **A.** Distribution of *Car5b/CA5B* expression in multiple tissues in both mouse (blue) and human (red). Expression levels are shown in TPM from the Expression Atlas of EMBL-EBL. **B.** Dynamic expression of *Car5b* during mouse development. Note that kidney shows the highest expression level of *Car5b* among tissues, which seems evident from embryonic stage ∼E17. **C.** Allelic expression analysis by BL6-specific ApaI digestion shows that *Car5b* does not escape in the mouse adult tissues tested. PCR products from amplification of *Car5b* cDNA from mouse adult brain, spleen, heart, kidney, liver and ovary, and from MEFs, all with the Xi from *spretus*, and control genomic DNA (gDNA) from BL6 and a F1 Bl6 x *spretus* heart were subject to mock or ApaI digestion (-/+) followed by gel electrophoresis. Complete digestion of PCR products from cDNA indicates the absence of transcripts from the *Car5b* allele on the *spretus* Xi, which lacks the ApaI site (see also additional file 1: Fig. S5). **D.** Allelic *Car5b* expression levels measured by allelic read counts and TPMs in mouse tissues and Patski cells confirm that *Car5b* escapes in Patski cells but not in brain, spleen, or ovary based on the ratio between Xi expression and total expression. RNA-seq data from Berletch et al., 2015 and Bonora et al., 2018.

**Figure S4.**
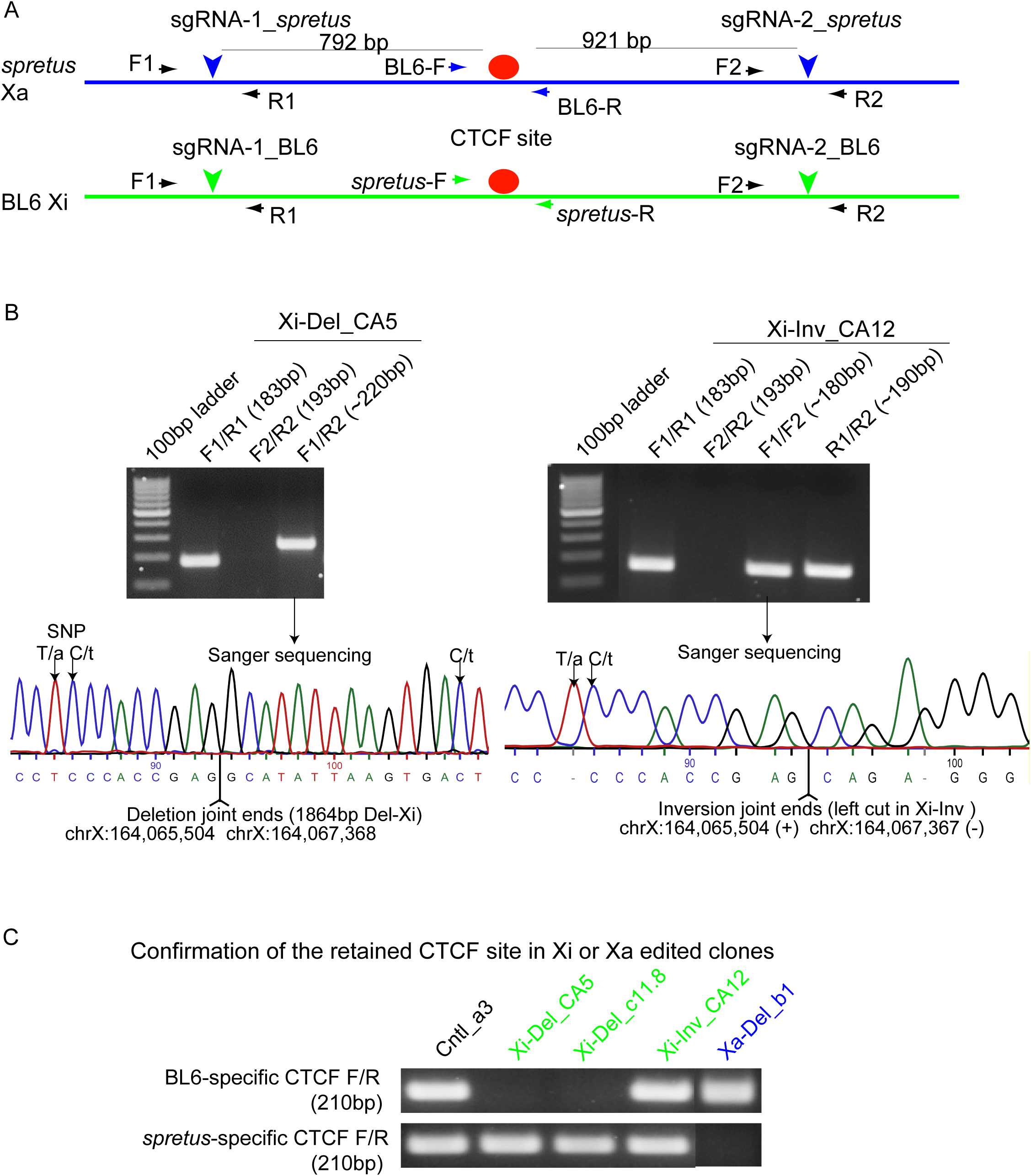
Allelic editing of CTCF site P4 at *Car5b*. **A.** Schematic of the ∼2kb CTCF binding region P4 upstream of *Car5b* (Fig. 2B). The positions of the BL6- and *spretus*-specific sgRNA pairs used for CRISPR/Cas9 editing of the Xa (blue) and Xi (green) are marked by downward arrowheads. The positions of the non-allelic (F1, R1, F2, R2) and allelic primers (F, R) are marked by black and colored arrows respectively. The CTCF site P4 is marked by a red oval. **B.** Single-cell clones with deletion or inversion of the ∼2kb region on the Xi or Xa were selected and verified by PCR using combinations of primers flanking the cutting sites (downward arrows in A) followed by Sanger sequencing to verify allelic editing. Examples of analysis of Xi-Del clone CA5 and Xi-Inv clone CA12 are shown. The SNP positions and the junction sequences are indicated on the Sanger sequence. **C.** PCR with allelic primers (F, R) flanking region P4 further confirms correct editing events demonstrating loss of the Xi-specific CTCF region P4 in Xi-Del, with retention in Xi-Inv and Xa-Del, while loss of the Xa-specific CTCF region P4 is lost in Xa-Del cells, and retained in Xi-Del and Xi-Inv.

**Figure S5.**
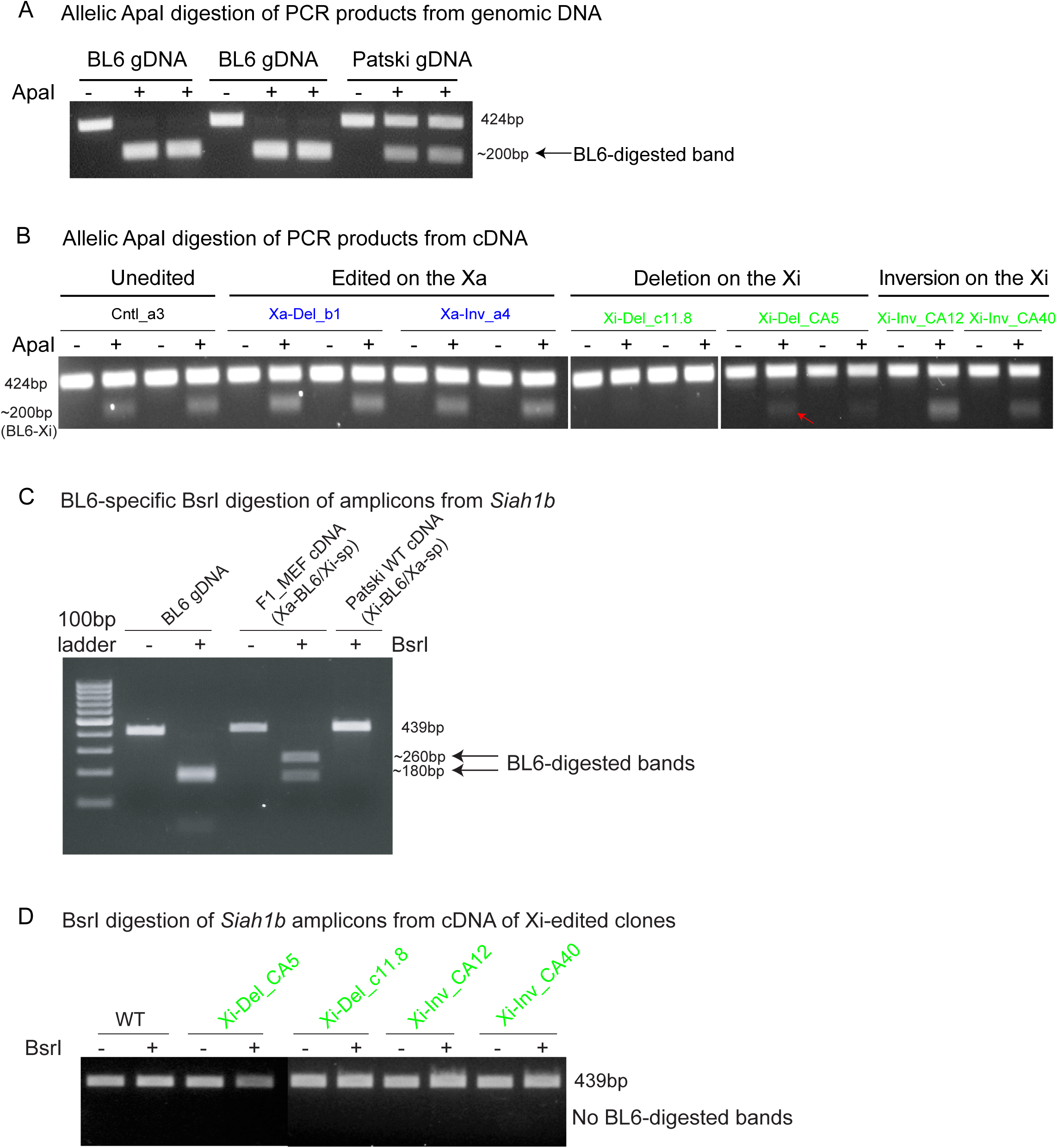
Allelic expression analysis using BL6-specific restriction enzyme digestion. **A-B.** BL6-specific ApaI digestion shows that only Xi-deletion of the CTCF P4 site abolishes *Car5b* escape in Patski cells. **A.** PCR products from genomic DNA from BL6 mice or Patski cells were subject to mock or ApaI digestion (-/+) followed by gel electrophoresis, which confirms BL6-specific ApaI digestion. **B.** PCR products from cDNA from CTCF P4 edited lines were subject to mock or ApaI digestion (-/+) followed by gel electrophoresis. Only lines with Xi-specific deletion of the CTCF site (Xi-Del) show the absence of a digested band, indicating loss of *Car5b* escape. Two biological replicates per line were tested except for two Xi-Inv lines. Note that in one of the two replicates of Xi-Del clone CA5, a very weak digested band (red arrow) was observed, indicating a much lower level of *Car5b* escape compared to the control (Ctrl-a3). This is consistent with the results from Q-PCR analysis in this line (Fig. 3C). **C-D**. BL6-specific BsrI digestion shows that editing of the CTCF region P4 located between *Car5b* and *Siah1b* on the Xi has no effects on *Siah1b*. **C.** PCR products from genomic DNA from BL6 mice, cDNAs from F1 hybrid MEFs and cDNA from Patski cells were subject to mock or ApaI digestion (-/+) followed by gel electrophoresis, which confirms BL6-specific BsrI digestion and no escape of *Siab1b* in either WT MEFs or Patski cells. **D.** PCR products from cDNA from Del-Xi lines were subject to mock or BsrI digestion (-/+) followed by gel electrophoresis. The absence of digested bands indicate absence of *Siab1b* escape.

**Figure S6.**
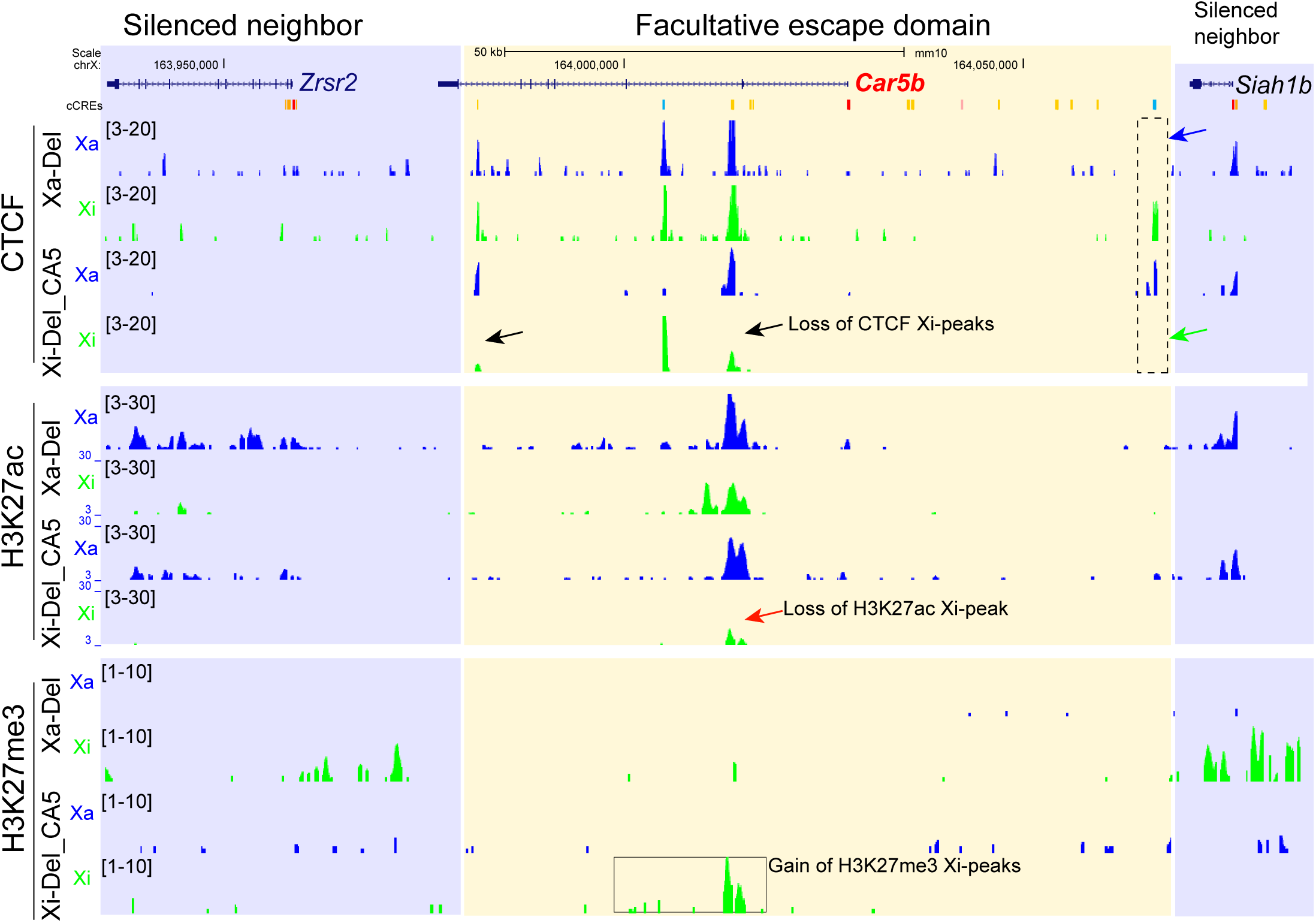
Confirmation of CTCF role in protecting the *Car5b* escape domain from changes in histone modifications. UCSC browser view of profiles of allelic CUT&RUN reads for CTCF, H3K27ac, and H3K27me3 are shown for the *Car5b* escape region in Xa-Del and in Xi-Del clone CA5. Similar but less pronounced effects than in clone 11.8 were observed, including loss of CTCF peaks within *Car5b*, loss of H3K27ac, and gain of H3K27me3 (Fig. 4A-C).

**Figure S7.**
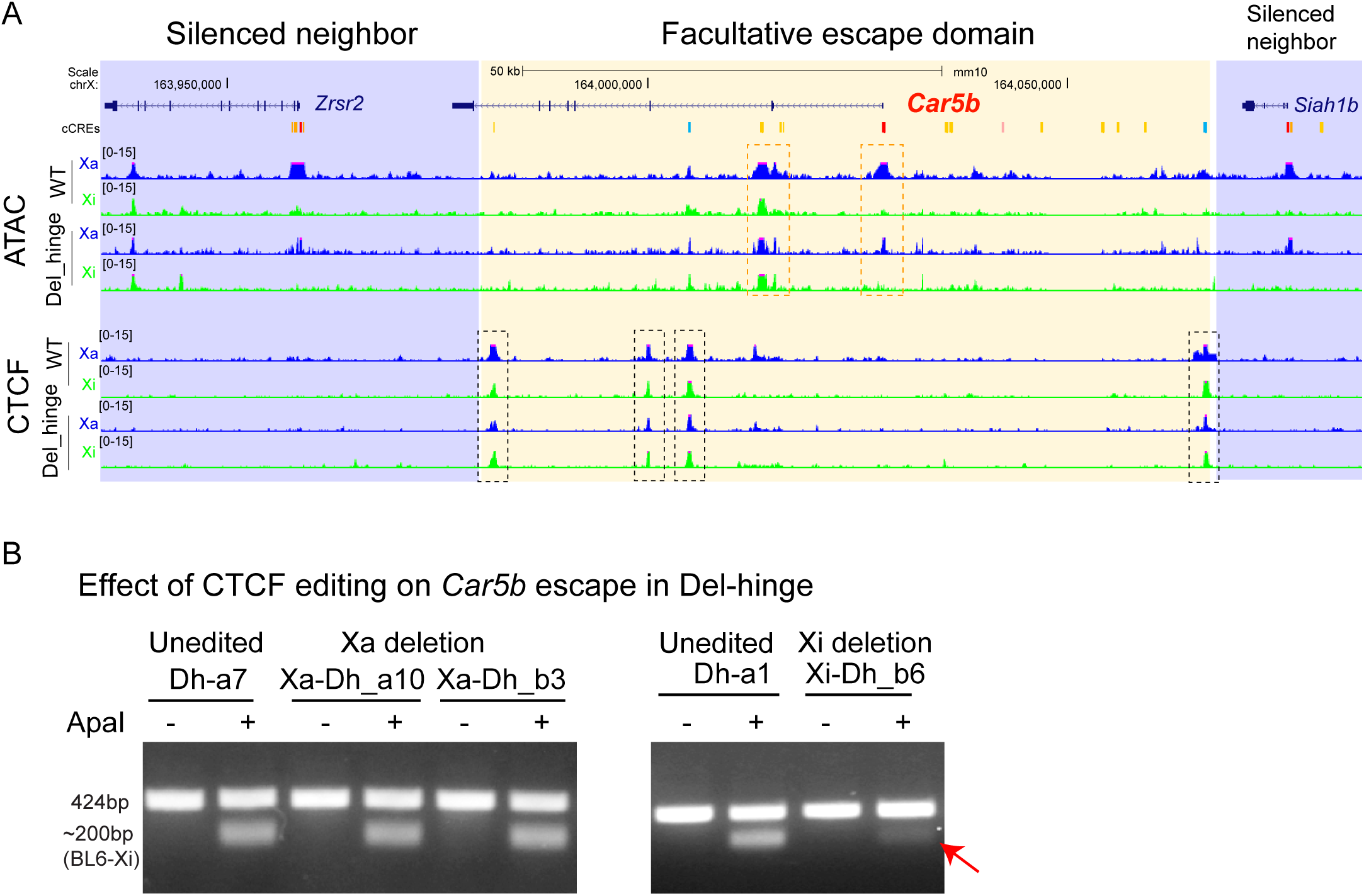
CTCF binding is required for *Car5b* escape in Patski cells with a *Dxz4* deletion. **A.** UCSC browser view of profiles of ATAC-seq reads and CTCF ChIP-seq reads on the Xa (blue) and Xi (green) are shown at *Car5b* in WT and Del-Hinge Patski cells. Relative to the profiles on the Xa allele in WT and Del-Hinge cells, only minor increases of ATAC-seq peaks at the promoter and enhancer (orange boxes) and of CTCF peaks (black boxes) are observed on the Xi allele in Del-Hinge cells, consistent a small increase of *Car5b* escape in Del-hinge (Fig. 5B). **B.** BL6-specific ApaI digestion shows that Xi-specific deletion of the CTCF P4 site strongly decreases *Car5b* escape in Patski cells with a *Dxz4* deletion. PCR products from cDNA from CTCF unedited and edited lines in Del-Hinge (Dh lines) were subject to mock or ApaI digestion (-/+) followed by gel electrophoresis. Xi-deletion of the CTCF site P4 in Del-Hinge (Dh-Xi_b6) shows a very weak digested band (red arrow), indicating a much reduced level of *Car5b* escape compared to the unedited or Xa-deleted lines.

**Figure S8.**
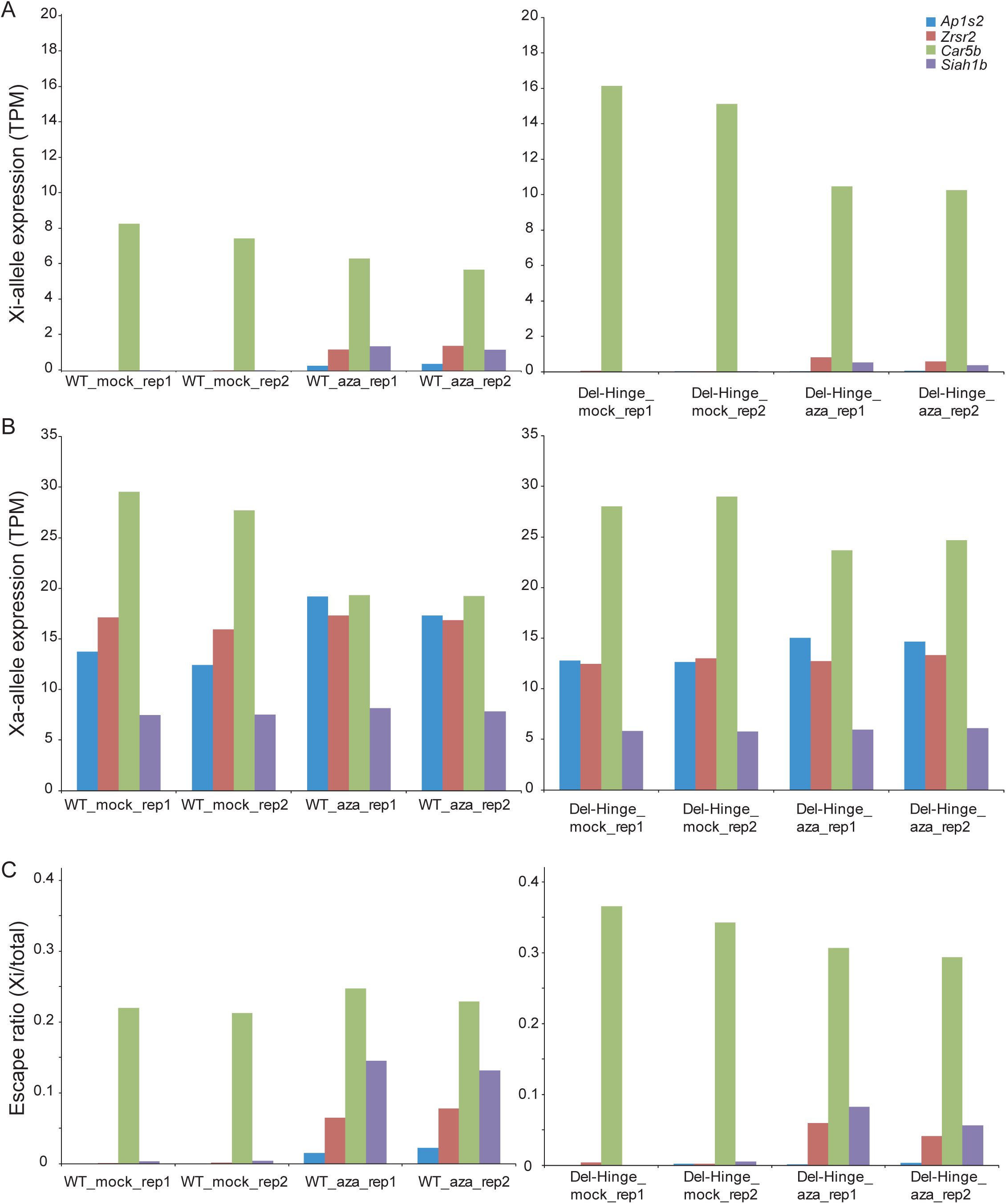
Inhibition of DNA methylation has no effect on *Car5b* escape but cause reactivation of neighbor genes subject to XCI. Allelic TPMs for the Xi allele, the Xa allele, and escape ratios (Xi expression versus total expression) were plotted for *Car5b* and neighbor genes *Ap1s2*, *Zrsf2*, and *Siah1b*. RNA-seq data from Bonora et al., 2018. Inhibition of DNA methylation was done by treatment of WT or Del-hinge Patski cells with DMSO (mock) or 4 µM 5-aza-2′-deoxycytidine (aza). Two biological replicates per treatment were done.

## Additional file 2

**Table S1. Information of escape domains on the X chromosome**

**Table S2. Information of motif analysis of CTCF Xi-peaks in Patski cells**

**Table S3. List of allelic sgRNAs for CRISP/Cas9 editing**

**Table S4. Information of primers used**

## References

1. Lyon MF. Gene action in the X-chromosome of the mouse (Mus musculus L.). Nature. 1961;190:372–3.

2. Loda A, Collombet S, Heard E. Gene regulation in time and space during X-chromosome inactivation. Nat Rev Mol Cell Biol. 2022;23:231–49.

3. Jacobson EC, Pandya-Jones A, Plath K. A lifelong duty: how Xist maintains the inactive X chromosome. Curr Opin Genet Dev. 2022;75:101927.

4. Fang H, Disteche CM, Berletch JB. X Inactivation and Escape: Epigenetic and Structural Features. Front Cell Dev Biol. 2019;7:219.

5. Posynick BJ, Brown CJ. Escape From X-Chromosome Inactivation: An Evolutionary Perspective. Front Cell Dev Biol. 2019;7:241.

6. Balaton BP, Cotton AM, Brown CJ. Derivation of consensus inactivation status for X-linked genes from genome-wide studies. Biol Sex Differ. England; 2015;6:35.

7. Balaton BP, Fornes O, Wasserman WW, Brown CJ. Cross-species examination of X-chromosome inactivation highlights domains of escape from silencing. Epigenetics Chromatin. England; 2021;14:12.

8. Berletch JB, Ma W, Yang F, Shendure J, Noble WS, Disteche CM, et al. Escape from X inactivation varies in mouse tissues. PLoS Genet. 2015;11:e1005079.

9. Tukiainen T, Villani A-C, Yen A, Rivas MA, Marshall JL, Satija R, et al. Landscape of X chromosome inactivation across human tissues. Nature. 2017;550:244–8.

10. Carrel L, Willard HF. X-inactivation profile reveals extensive variability in X-linked gene expression in females. Nature. England; 2005;434:400–4.

11. Filippova GN, Cheng MK, Moore JM, Truong J-P, Hu YJ, Nguyen DK, et al. Boundaries between chromosomal domains of X inactivation and escape bind CTCF and lack CpG methylation during early development. Dev Cell. 2005;8:31–42.

12. Horvath LM, Li N, Carrel L. Deletion of an X-inactivation boundary disrupts adjacent gene silencing. PLoS Genet. 2013;9:e1003952.

13. Peeters SB, Korecki AJ, Simpson EM, Brown CJ. Human cis-acting elements regulating escape from X-chromosome inactivation function in mouse. Hum Mol Genet. 2018;27:1252–62.

14. Qian J, Guan X, Xie B, Xu C, Niu J, Tang X, et al. Multiplex epigenome editing of MECP2 to rescue Rett syndrome neurons. Sci Transl Med. 2023;15:eadd4666.

15. Bonora G, Deng X, Fang H, Ramani V, Qiu R, Berletch JB, et al. Orientation-dependent Dxz4 contacts shape the 3D structure of the inactive X chromosome. Nat Commun. 2018;9:1445.

16. Nanni L, Ceri S, Logie C. Spatial patterns of CTCF sites define the anatomy of TADs and their boundaries. Genome Biol. 2020;21:197.

17. Ziebarth JD, Bhattacharya A, Cui Y. CTCFBSDB 2.0: a database for CTCF-binding sites and genome organization. Nucleic Acids Res. 2013;41:D188–194.

18. ENCODE Project Consortium, Moore JE, Purcaro MJ, Pratt HE, Epstein CB, Shoresh N, et al. Expanded encyclopaedias of DNA elements in the human and mouse genomes. Nature. 2020;583:699–710.

19. Keown CL, Berletch JB, Castanon R, Nery JR, Disteche CM, Ecker JR, et al. Allele-specific non-CG DNA methylation marks domains of active chromatin in female mouse brain. Proc Natl Acad Sci U S A. 2017;114:E2882–90.

20. Tian D, Sun S, Lee JT. The long noncoding RNA, Jpx, is a molecular switch for X chromosome inactivation. Cell. 2010;143:390–403.

21. Chureau C, Chantalat S, Romito A, Galvani A, Duret L, Avner P, et al. Ftx is a non-coding RNA which affects Xist expression and chromatin structure within the X-inactivation center region. Hum Mol Genet. 2011;20:705–18.

22. Lopes AM, Arnold-Croop SE, Amorim A, Carrel L. Clustered transcripts that escape X inactivation at mouse XqD. Mamm Genome Off J Int Mamm Genome Soc. 2011;22:572–82.

23. Yang F, Babak T, Shendure J, Disteche CM. Global survey of escape from X inactivation by RNA-sequencing in mouse. Genome Res. 2010;20:614–22.

24. Cotton AM, Ge B, Light N, Adoue V, Pastinen T, Brown CJ. Analysis of expressed SNPs identifies variable extents of expression from the human inactive X chromosome. Genome Biol. 2013;14:R122.

25. Weintraub AS, Li CH, Zamudio AV, Sigova AA, Hannett NM, Day DS, et al. YY1 Is a Structural Regulator of Enhancer-Promoter Loops. Cell. 2017;171:1573–1588.e28.

26. Skene PJ, Henikoff S. An efficient targeted nuclease strategy for high-resolution mapping of DNA binding sites. eLife. 2017;6:e21856.

27. Fang H, Bonora G, Lewandowski JP, Thakur J, Filippova GN, Henikoff S, et al. Trans- and cis-acting effects of Firre on epigenetic features of the inactive X chromosome. Nat Commun. 2020;11:6053.

28. Vallender TW, Lahn BT. Improved DNA methylation analysis via enrichment of demethylated cells expressing an X-inactivated transgene. BioTechniques. 2006;41:461–6.

29. Chen X, Ke Y, Wu K, Zhao H, Sun Y, Gao L, et al. Key role for CTCF in establishing chromatin structure in human embryos. Nature. 2019;576:306–10.

30. Dehingia B, Milewska M, Janowski M, Pękowska A. CTCF shapes chromatin structure and gene expression in health and disease. EMBO Rep. 2022;23:e55146.

31. Popay TM, Dixon JR. Coming full circle: On the origin and evolution of the looping model for enhancer-promoter communication. J Biol Chem. 2022;298:102117.

32. Chen C-Y, Shi W, Balaton BP, Matthews AM, Li Y, Arenillas DJ, et al. YY1 binding association with sex-biased transcription revealed through X-linked transcript levels and allelic binding analyses. Sci Rep. England; 2016;6:37324.

33. Tang Z, Luo OJ, Li X, Zheng M, Zhu JJ, Szalaj P, et al. CTCF-Mediated Human 3D Genome Architecture Reveals Chromatin Topology for Transcription. Cell. 2015;163:1611–27.

34. Deng X, Ma W, Ramani V, Hill A, Yang F, Ay F, et al. Bipartite structure of the inactive mouse X chromosome. Genome Biol. 2015;16:152.

35. Splinter E, de Wit E, Nora EP, Klous P, van de Werken HJG, Zhu Y, et al. The inactive X chromosome adopts a unique three-dimensional conformation that is dependent on Xist RNA. Genes Dev. 2011;25:1371–83.

36. Yu B, Qi Y, Li R, Shi Q, Satpathy AT, Chang HY. B cell-specific XIST complex enforces X-inactivation and restrains atypical B cells. Cell. 2021;184:1790–1803.e17.

37. Yang T, Ou J, Yildirim E. Xist exerts gene-specific silencing during XCI maintenance and impacts lineage-specific cell differentiation and proliferation during hematopoiesis. Nat Commun. 2022;13:4464.

38. Giorgetti L, Lajoie BR, Carter AC, Attia M, Zhan Y, Xu J, et al. Structural organization of the inactive X chromosome in the mouse. Nature. 2016;535:575–9.

39. Lingenfelter PA, Adler DA, Poslinski D, Thomas S, Elliott RW, Chapman VM, et al. Escape from X inactivation of Smcx is preceded by silencing during mouse development. Nat Genet. 1998;18:212–3.

40. Thakur J, Fang H, Llagas T, Disteche CM, Henikoff S. Architectural RNA is required for heterochromatin organization. bioRxiv. 2019;784835.

